# Transient inhibition of translation improves long-term cardiac function after ischemia/reperfusion by attenuating the inflammatory response

**DOI:** 10.1101/2022.07.25.501397

**Authors:** Christoph Hofmann, Adrian Serafin, Ole M Schwerdt, Fereshteh S Younesi, Florian Sicklinger, Ingmar Sören Meyer, Ellen Malovrh, Clara Sandmann, Lonny Jürgensen, Verena Kamuf-Schenk, Claudia Stroh, Zoe Löwenthal, Mandy Rettel, Frank Stein, Hugo A. Katus, Tobias Jakobi, Norbert Frey, Florian Leuschner, Mirko Völkers

**Author notes:** Correspondence to: Mirko Völkers, University Hospital Heidelberg, Department of Cardiology, Angiology, and Pneumology, Internal Medicine III, Im Neuenheimer Feld 410, 69120 Heidelberg, Germany.

## Abstract

**Rationale:** Rapid reperfusion is the most effective treatment for attenuating cardiac injury caused by myocardial ischemia. Yet, reperfusion itself elicits damage to the myocardium through incompletely understood mechanisms, known as ischemia/reperfusion (I/R) injury. The myocardium adapts to I/R by changes in gene expression, which determines the cellular response to reperfusion. Protein translation is a key component of gene expression. However, it is unknown how regulation of translation contributes to cardiac gene expression in response to reperfusion and whether it can be targeted to mitigate I/R injury.

**Methods:** To examine translation and its impact on gene expression in response to I/R we assessed protein synthesis at different timepoints after ischemia and reperfusion in vitro and in vivo. Pharmacological inhibitors were used to dissect the underlying molecular mechanisms of translational control. Transient inhibition of protein synthesis was undertaken to decipher the effects of the translational response to reperfusion on cardiac function and inflammation. Cell-type-specific ribosome profiling was performed in mice subjected to I/R to determine the impact of translation on the regulation of gene expression in cardiomyocytes.

**Results:** Reperfusion increased translation rates from a previously suppressed state during ischemia in cardiomyocytes, which was associated with the induction of cell death. In vivo, I/R resulted in strong activation of translation in the myocardial border zone. Detailed analysis revealed that the upregulation of translation is mediated by eIF4F complex formation, which was specifically mediated by the mTORC1-4EBP1-eIF4F axis. Short-term pharmacological inhibition of eIF4F complex formation by 4EGI-1 or rapamycin, respectively, attenuated translation, reduced infarct size and improved long-term cardiac function after myocardial infarction. Cardiomyocyte-specific ribosome profiling identified that reperfusion damage increased translation of mRNA networks in cardiomyocytes associated with cardiac inflammation and cell infiltration. Transient inhibition of the mTORC1-4EBP1-eIF4F axis decreased the expression of proinflammatory transcripts such as Ccl2, thereby reducing Ly6C^hi^ monocyte infiltration and myocardial inflammation.

**Conclusions:** Myocardial reperfusion induces protein synthesis in the border zone which contributes to I/R injury by rapidly translating a specific maladaptive mRNA network that mediates immune cell infiltration and inflammation. Transient inhibition of the mTORC1-4EBP1-eIF4F signaling axis during reperfusion attenuates this proinflammatory translational response, protects against I/R injury and improves long-term cardiac function after myocardial infarction.

**Clinical Perspective:** *What Is New?:*

- This is the first study to investigate the impact of translational regulation on cardiomyocyte gene expression in response to myocardial ischemia/reperfusion.
- We show that translation regulates approximately two-thirds of differentially expressed genes in cardiomyocytes after ischemia/reperfusion, including many involved in inflammation and immune cell infiltration.
- The translational response to ischemia/reperfusion is regulated by the mTORC1-4EBP1-eIF4F axis, which determines pro-inflammatory monocyte infiltration via control of the expression of the chemokine Ccl2.

*What Are the Clinical Implications?:*

- Currently, there are no specific therapies to prevent ischemia/reperfusion injury, which is mediated, at least in part, by a maladaptive inflammatory response.
- A translationally controlled network regulated by the mTORC1-4EBP1-eIF4F axis can be targeted by a short-term pharmacological intervention to attenuate the inflammatory response and improve cardiac function after ischemia/reperfusion in mice.
- This study supports the emerging concept of selectively inhibiting maladaptive elements of the inflammatory response to improve outcome in patients after myocardial infarction; in addition, it provides a mechanistic basis for the currently ongoing CLEVER-ACS trial.

## Introduction

Myocardial infarction is a leading cause of death worldwide.^1^ It is caused by inadequate blood supply to the myocardium, resulting in ischemia and subsequent acute cardiac injury. Prolonged duration of ischemia causes progressive and irreversible loss of viable cardiac tissue. Therefore, the most effective treatment for mitigating acute myocardial injury is rapid reperfusion, which restores cellular oxygen and nutrients and thereby prevents further cell death resulting from ischemia.^2^ Timely reperfusion decreases cardiac morbidity and mortality and is the primary therapeutic strategy for treating acute myocardial infarction.^2^ However, reperfusion itself causes myocardial injury by still incomplete understood mechanisms referred to as I/R injury.^3^ Despite the continuously improving success of reperfusion therapy for acute myocardial infarction,^1^ therapies for myocardial infarction currently target only the ischemic aspect, whereas no intervention to reduce reperfusion injury has been successfully translated into the clinic.^3^ Thus, there is great need to define novel mechanisms of reperfusion injury that could be targeted for the treatment of acute myocardial infarction.

In response to stress, cells react with rapid changes in gene expression. Due to technical limitations, gene expression has been studied mainly by mRNA sequencing, however the correlation between transcript and protein levels is frequently poor.^4–6^ Recent studies could demonstrate that translation plays a fundamental role in regulating gene expression in the diseased heart.^5–9^ However, how translation is regulated in response to myocardial reperfusion and to what extend this contributes to the regulation of gene expression is largely unknown.

Translation is primarily regulated at the initiation stage, which is the assembly of a ribosome on the initiation codon of an mRNAs open reading frame.^10^ For the majority of transcripts, mRNAs selected for translation initiation are activated by the binding of the eukaryotic initiation factor (eIF) 4F, a heterotrimeric protein complex, consisting of eIF4E, eIF4G and eIF4A that binds the 5’ cap of mRNAs to promote translation. This process is controlled by mechanistic target of rapamycin complex 1 (mTORC1), a cellular hub that integrates information about the availability of nutrients, growth factors, energy, oxygen and stress to coordinate translation through its kinase activity toward eIF4E-binding protein 1 (4EBP1). Phosphorylation of 4EBP1 prevents its inhibitory binding to eIF4E, freeing eIF4E to bind eIF4G and participate in translation initiation. Previous studies have shown that inhibition of mTORC1 protects against ischemic damage and I/R injury,^11–14^ which is currently being investigated in a randomized, prospective, double-blind, multicenter phase I-II clinical trial (CLEVER-ACS; NCT01529554).^15^ However, the mechanistic basis of the cardioprotective effect of pharmacological mTORC1 inhibition and whether it is mediated by translational regulation of gene expression remain to be fully elucidated.

Here, we present a detailed analysis of the translational response of cardiomyocytes to myocardial ischemia and reperfusion. Our analysis reveals extensive regulation of protein synthesis and gene expression by translation in response to reperfusion, mediated at least in part by activation of the mTORC1-4EBP1-eIF4F axis in border zone cardiomyocytes. By combining ribosome profiling (Ribo-Seq) with a cardiomyocyte-specific ribosome tagging approach in mice, we identified a translationally controlled maladaptive mRNA network in cardiomyocytes that mediates proinflammatory Ly6C^hi^ monocyte infiltration and inflammation. Transient inhibition of mTORC1-dependent translation during reperfusion reduced proinflammatory cell infiltration, decreased infarct size, and improved long-term cardiac function after myocardial infarction. Overall, our data demonstrate a maladaptive, translationally regulated inflammatory response of border zone cardiomyocytes after reperfusion that can be targeted by a transient pharmacological intervention to improve long-term cardiac function after myocardial infarction.

## Methods

All animal experiments were approved by the institutional animal care and use committee of Heidelberg University. Data, analytical methods, and study materials are available from the corresponding author on reasonable request. Raw sequencing data have been made publicly available at the SRA and can be accessed under PRJNA830657. The mass spectrometry proteomics data have been deposited to the ProteomeXchange Consortium via the PRIDE^16^ partner repository with the dataset identifier PXD035154. Additional details of experimental procedures are provided in the Data Supplement.

## Results

### Activation of translation in response to ischemia/reperfusion

To examine the translational response of cardiomyocytes subjected to increasing periods of simulated ischemia (sI) or sI for 3 hours followed by reperfusion (sI/R), we labeled newly synthesized proteins with puromycin in vitro in isolated neonatal rat cardiomyocytes (NRCMs).^17^ As expected, this revealed a progressive downregulation of translation in response to sI (Figure 1A). In contrast, reperfusion increased translation, resulting in rapid restoration of baseline protein synthesis rates within 3 hours (Figure 1B). To validate these results in vivo, mice were subjected to permanent ligation of the left anterior descending artery (LAD) ligation or I/R surgery followed by i.p. injection of puromycin shortly before isolation of the hearts. Similar to NRCMs, ischemia attenuated myocardial translation after 3 hours of LAD ligation (Figure 1C and 1D). The reduction of protein synthesis rates was negatively correlated with plasma CK levels, suggesting that the extend of translational repression is associated with the infarct size (Figure 1E). In line with the in vitro data, translation was significantly upregulated 2 days after reperfusion (Figure 1F and G), which was positively correlated with plasma troponin T levels (Figure 1H), suggesting that a larger infarcts size results in an increased translational response to I/R.

**Figure 1.**
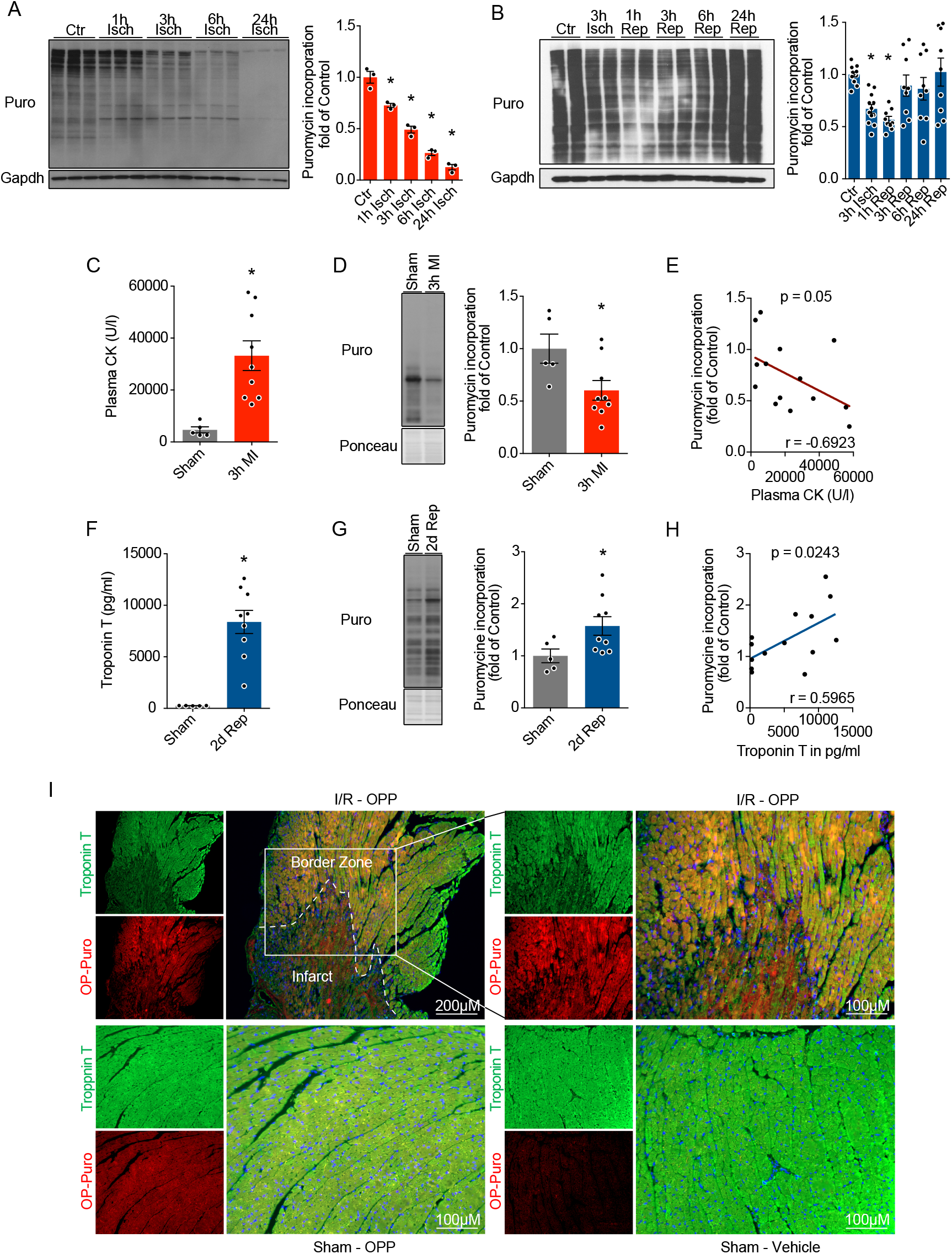
Dynamic regulation of myocardial translation rates in response to ischemia and reperfusion. **A** and **B**, Puromycin immunoblots with quantification of NRCMs at increasing timepoints of simulated ischemia, n = 3 (**A**) or ischemia (3 hours) followed by reperfusion, n = 9-12 (**B**). **C** to **E**, Plasma CK levels, n = 5-9 (**C**), puromycin incorporation, measured by immunoblot, n = 5-9 (**D**) and correlation of Plasma CK and cardiac puromycin incorporation (**E**) of mice 3 hours after Sham surgery or LAD ligation (MI). **F** to **I**, Plasma Troponin T levels, n = 5-9 (**F**), puromycin incorporation, measured by immunoblot, n = 5-9 (**G**), correlation of Plasma Troponin T and cardiac puromycin incorporation (**H**) and OP-Puromycin immunostaining of the infarct region and border zone (**I**) of mice 2 days after Sham or I/R surgery. Ctr (control), Isch (ischemia), Rep (reperfusion), MI (myocardial infarction), I/R (ischemia/reperfusion surgery), Puro (puromycin), * indicates p<0.05 from control or sham. For statistical analysis one-way ANOVA with Turkey post-hoc analysis was used for **A** and **B**. An unpaired two tailed t-test was used for **C, D, F** and **G**. A Pearson correlation with two-tailed p value was computed with GraphPad Prism 7.0 for **E** and **H**. p < 0.05 was defined as significant difference. Error bars show standard error of the mean (SEM).

To further characterize the translational response after I/R in vivo, we performed immunofluorescent imaging of mouse heart sections 2 days after reperfusion. Mice were treated with O-propargyl-puromycin (OP-puromycin), an alkyne analog of puromycin with improved pharmacological properties for visualization of protein synthesis in vivo.^18^ Incorporation of OP-puromycin in response to reperfusion was predominantly observed in the border zone (Figure 1I). In conclusion, myocardial reperfusion results in the activation of translation from a previously suppressed state during ischemia, which appears to be mediated in a significant extend by border zone cells, including cardiomyocytes.

### The translational response after reperfusion is mediated by the mTORC1-4EBP1-eIF4F axis

Translation rates are controlled at the level of initiation, which is regulated in large part by the binding of eIF4F to the 5’-cap of mRNA (Figure 2A). To quantify the regulation of translation initiation by eIF4F-mRNA complex formation during sI/R in NRCMs, we performed immunoprecipitation of mRNA cap-binding proteins via 7-methylguanosin (m^7^GTP)-coupled agarose beads. Ischemia induced the binding of 4EBP1 to the cap-mimic, which inhibited eIF4F complex formation and resulted in decreased association of eIF4G and eIF4A with eIF4E (Figure 2B). This was completely reversed by reperfusion at 6h, resulting in complete restoration of eIF4F assembly (Figure 2B). To systematically identify the upstream signaling events that control eIF4F-dependent translation initiation, we next characterized the activity of key signaling pathways in NRCMs at different timepoints of ischemia and reperfusion that are thought to regulate eIF4F complex activity. eIF4F-depentent translation is primarily controlled by the mTORC1-4EBP1-eIF4F pathway (Figure 2A).^19^ In addition, it was previously proposed that the association of mRNAs with eIF4F is regulated by phosphorylation of eIF4E at S209 via mitogen-activated protein kinase-interacting kinases 1 and 2 (Mnk1/2) (Figure 2A),^20,21^ which is mediated by ERK1/2 in NRCMs (Online Figure 1). Both the mTORC1-4EBP1-eIF4F and ERK1/2-MNK1/2-eIF4E^S209^ pathways were strongly regulated in response to sI/R and were therefore considered potential mediators of eIF4F-dependent translational regulation (Figure 2C and 2D).

**Figure 2.**
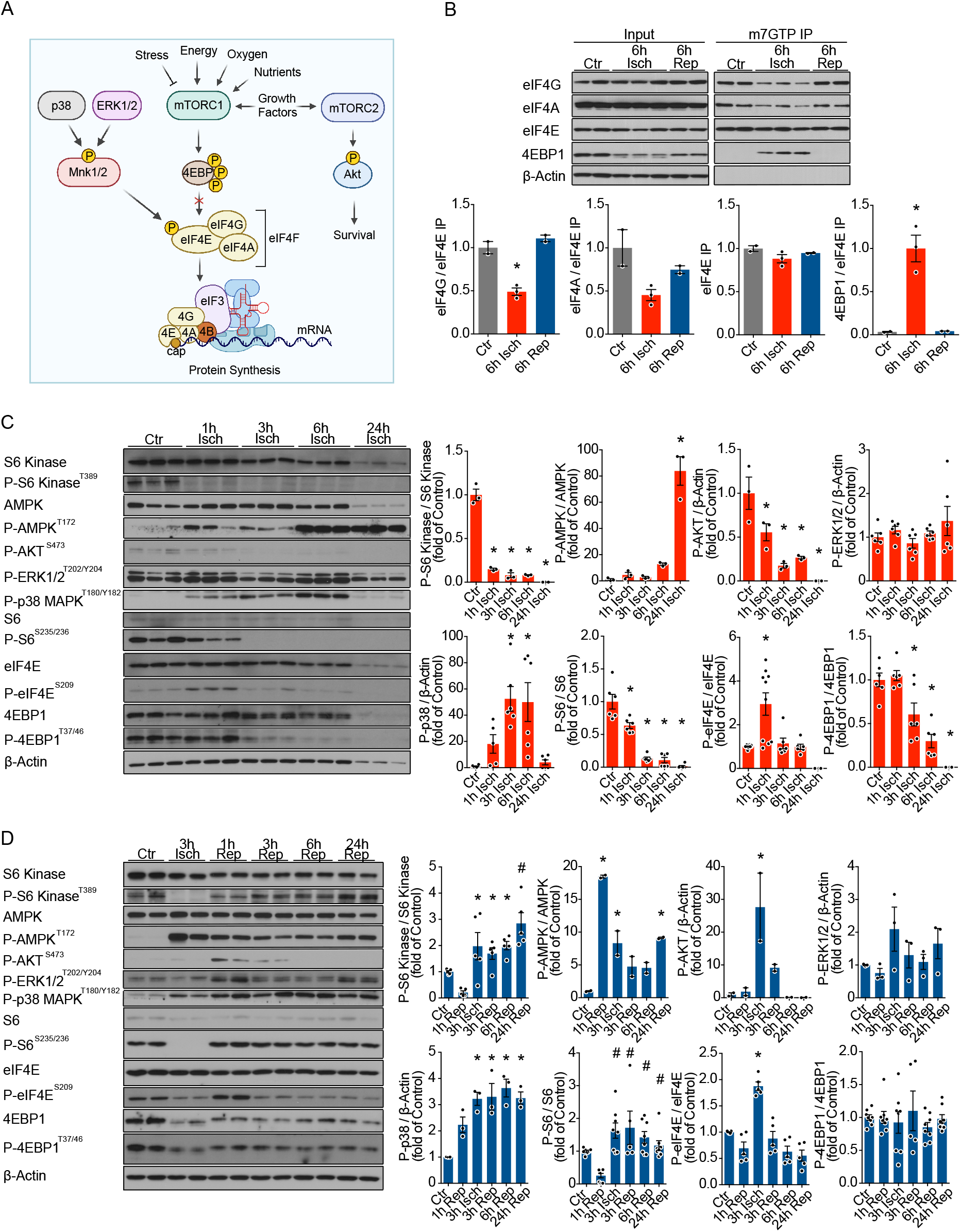
Activation of signaling pathways controlling eIF4F-dependent translation initiation after reperfusion. **A**, Diagram of major signaling pathways proposed to regulate eIF4F-dependent translation initiation. **B**, Quantification of eIF4F complex assembly by immunoprecipitation of mRNA cap-binding proteins via m^7^GTP-coupled agarose beads after simulated ischemia or ischemia (6 hours) followed by reperfusion in NRCMs, n = 2-3. **C** and **D**, Immunoblots and quantifications of proposed eIF4F-regulatory pathways of NRCMs at increasing timepoints of simulated ischemia, 3-10 (**C**) or ischemia (3 hours) followed by reperfusion, n = 2-8 (**D**). Ctr (control), Isch (ischemia), Rep (reperfusion), * indicates p<0.05 from control. # indicates p<0.05 from 3h ischemia. For statistical analysis one-way ANOVA with Turkey post-hoc analysis was used for **B, C** and **D**. p < 0.05 was defined as significant difference. Error bars show standard error of the mean (SEM). (A) made in ©BioRender - biorender.com.

We next sought to determine the extent to which each signaling pathway affects eIF4F-cap interaction and translation rates in response to reperfusion. We used the mTORC1 inhibitor rapamycin (sirolimus), the ATP-competitive mTOR inhibitor torin1, the Mnk1/2 inhibitor eFT508^22^ and the eIF4F-competitive eIF4E/eIF4G interaction inhibitor 4EGI-1^23^, to specifically target each of the sub-pathways thought to be involved in eIF4F-dependent translation initiation (Figure 3A). For each inhibitor, we tested the minimal effective dose required to sufficiently inhibit its regulated signaling cascade in NRCMs (Figure 3B). High doses of 4EGI-1 have previously been described as toxic to cardiomyocytes,^24^ consistent with the apoptotic potential of several other compounds interacting with eIF4E.^23,25–27^ We confirmed this observation for high doses, however, a concentration of 25μM to 100μM of 4EGI-1 was sufficient to significantly inhibit eIF4F complex formation and translation in NRCMs without causing cell death within 24 hours (Online Figure 2).

**Figure 3.**
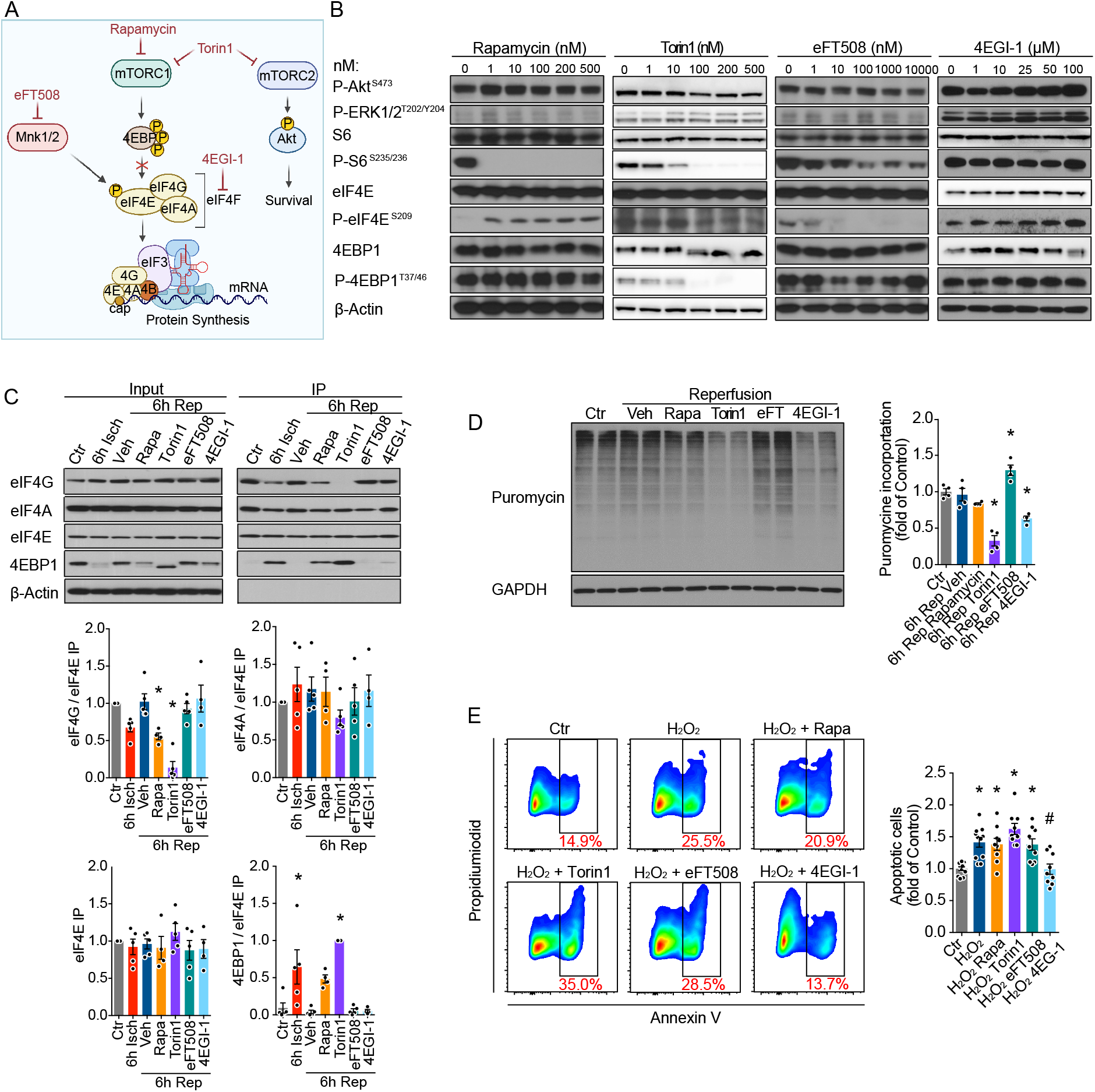
Control of translation rates in response to reperfusion by the mTORC1-4EBP1-eIF4F signaling axis. **A**, Diagram of signaling pathways affected by the pharmacological compounds used to modulate translation initiation in this study. **B**, Representative immunoblots of the activity of eIF4F-regulatory pathways in response to increasing doses of the indicated pharmacological compounds in NRCMs. **C** and **D**, Immunoprecipitation (IP) of eIF4F proteins via m^7^GTP-coupled agarose beads, n = 4-5 (**C**) and puromycin immunoblot, n = 4 (**D**) of NRCMs in response to sI/R and pharmacological inhibitor treatment. A representative immunoblot is shown on the left, and quantification of all replicates is shown on the right **E**, Quantification of NRCM apoptosis by FACS analysis 4 hours after simultaneous H_2_O_2_ (50 μM) and pharmacological inhibitor treatment, n = 9-12. Representative FACS plots of each condition are shown on the left. The following concentrations were used for **C, D** and **E**: Rapamycin 100nM, Torin1 100nM, eFT508 100nM, 4EGI-1 100μM. Ctr (control), Isch (ischemia), Rep (reperfusion), * indicates p<0.05 from control. # indicates p<0.05 from H_2_O_2_. For statistical analysis one-way ANOVA with Turkey post-hoc analysis was used for **C, D** and **E**. p < 0.05 was defined as significant difference. Error bars show standard error of the mean (SEM). (A) made in ©BioRender - biorender.com.

All pharmacological mTORC1-4EBP1-eIF4F axis inhibitors tested prevented eIF4F complex formation by reducing the dissociation of 4EBP1 from eIF4E in response to reperfusion (Figure 3C and Online Figure 2). This was associated with significant inhibition of puromycin incorporation, confirming the involvement of this pathway in translational upregulation after reperfusion in NRCMs (Figure 3D and Online Figure 3). Importantly, treatment of NRCMs with the eIF4E/eIF4G interaction inhibitor 4EGI-1 protected from apoptosis in an in vitro model of reperfusion injury by H_2_O_2_, indicating that this pathway is involved in the regulation of cell death in response to reperfusion (Figure 3E).

In contrast, inhibition of the MNK1/2-eIF4E^S209^ axis by eFT508 had no effect on eIF4F-cap interaction, protein synthesis rates, or apoptosis (Figure 3C to 3E). To confirm that eIF4E^S209^ phosphorylation is not involved in regulating translation rates we generated S209-phospho-dead-eIF4E (mutant-eIF4E). Neither overexpression of WT-eIF4E nor mutant-eIF4E affected translation rates (Online Figure 4). Moreover, the MNK1/2 inhibitor eFT508 had no effect on plasma TnT levels or cardiac function in a small series of mice after I/R surgery in vivo (Online Figure 4). Thus, we decided to focus our subsequent experiments primarily on the mTORC1-4EBP1-eIF4F pathway.

**Figure 4.**
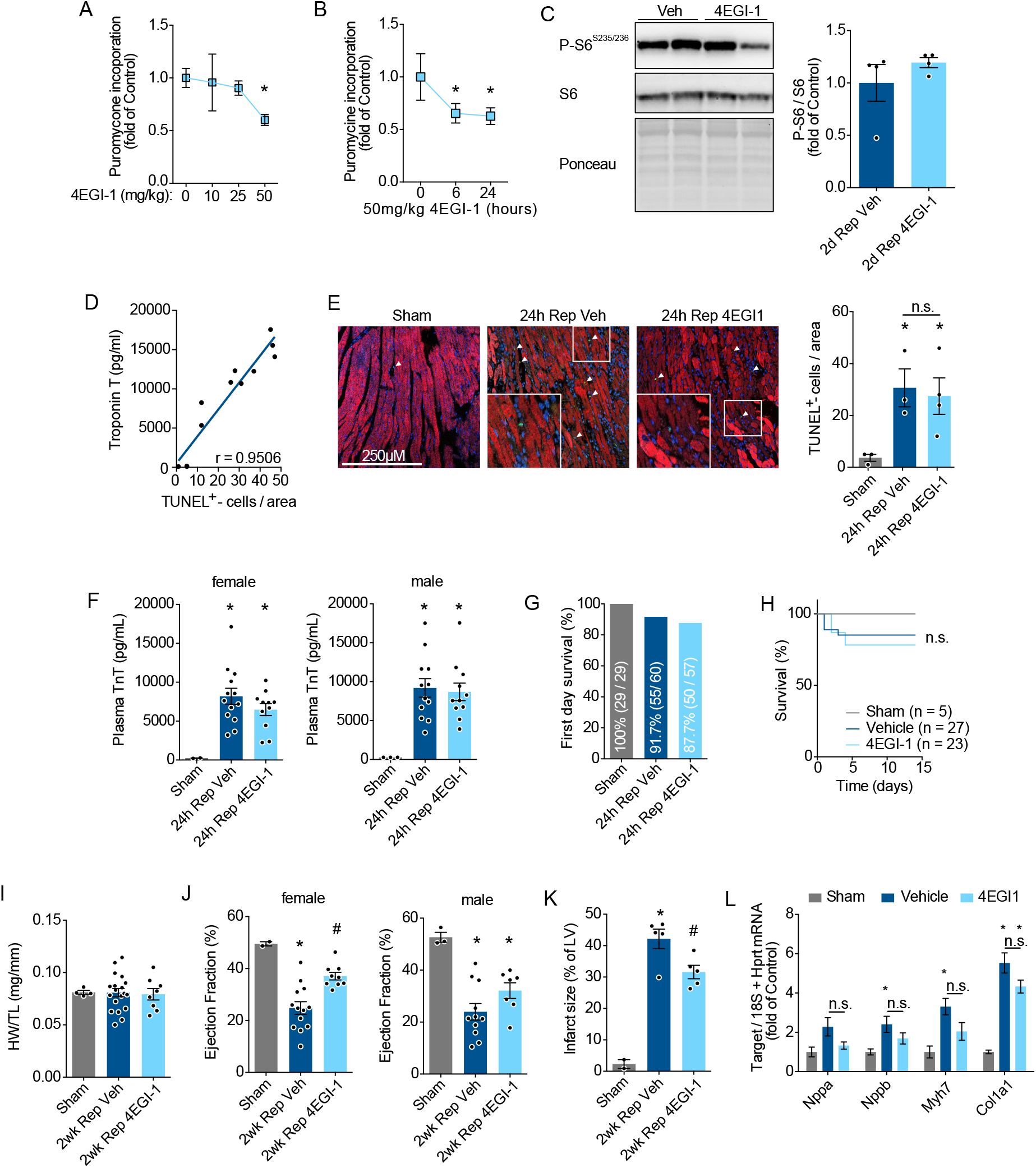
Inhibition of translation during reperfusion protects against I/R injury and improves long-term cardiac function. **A** and **B**, Quantification of myocardial puromycin incorporation in vivo after 6 hours of 4EGI-1 treatment with different concentrations, n = 6 (0, 25 and 50 mg/kg); n = 2 (10 mg/kg), (**A**) or at different timepoints after 4EGI-1 treatment, n = 5 (control), n = 4 (6 hours), n = 3 (24 hours) (**B**) as assessed by immunoblot. Puromycin was i.p. injected 30 minutes before animals were sacrificed. **C**, S6^S235/236^ phosphorylation of left ventricular lysates of vehicle or 50 mg/kg 4EGI-1 treated mice 2 days after I/R surgery assessed by immunoblot, n = 4. **D** and **E**, Correlation of plasma Troponin T levels and TUNEL^+^-cells (**D**) and representative immunostaining and quantification of TUNEL^+^-cells per view by immunofluorescence, n = 3-4 (**E**) in sham or I/R surgery operated mice treated with vehicle or 50 mg/kg 4EGI-1. **F**, Plasma Troponin T levels at 24 hours after I/R surgery in female (n = 2-13) and male mice (n = 3-12). **G**, 24-hour survival of all sham or I/R surgery operated mice after vehicle or 50 mg/kg 4EGI-1 treatment used in this study, n = 29-60. **H**, 2-week survival of sham or I/R surgery operated mice treated with vehicle or 50 mg/kg 4EGI-1 corresponding to mice used for I to L, n = 5-27. **I, J, K** and **L**, Heart weight (HW) to body weight (BW) ratio (**I**), left ventricular ejection fraction of female (n = 2-13) and male mice (n = 3-11) (**J**), infarct size (n = 2-5, male mice) (**K**), and left ventricular expression of *Nppa, Nppb, Myh7* and *Col1a1* (n = 4 for sham, n = 6 for vehicle and 4EGI-1, female mice) (**L**) 2 weeks after sham or I/R surgery operated mice treated with vehicle or 50 mg/kg 4EGI-1. Rep (reperfusion), Veh (vehicle). * indicates p<0.05 from control or sham. # indicates p<0.05 from 2wk Rep Veh. For statistical analysis one-way ANOVA with Turkey post-hoc analysis was used for **A, B, E, F, I, J, K** and **L**. An unpaired two tailed t-test was used for **C**. A survival curve analysis was performed with GraphPad Prism 7.0 for **H**. p < 0.05 was defined as significant difference. Error bars show standard error of the mean (SEM).

### Selective inhibition of mTORC1-dependent translation during reperfusion improves myocardial function and reduces infarct size after cardiac ischemia/reperfusion

We identified the mTORC1-4EBP1-eIF4F pathway as a master regulator of translation rates in response to sI/R in NRCMs (Figure 2 and 3). Previous studies have consistently demonstrated upregulation of mTORC1 activity after I/R in vivo,^14,28,29^ which we could reproduce (Online Figure 5). Similar to the observation of activated translation in the border zone, mTORC1 appeared to be activated in the myocardial border zone 2 days after reperfusion (Online Figure 5). To further study the functional relevance of mTORC1-dependent translational control after reperfusion, we examined a protocol of short-term inhibition of the mTORC1 pathway with the clinically approved first-generation mTOR inhibitor rapamycin, which we based on the protocol of the currently ongoing CLEVER-ACS trial (NCT01529554, initial loading dose followed by a short-term maintenance dose).^15^ This could overcome maladaptive off-target effects toward mTORC2 that occurs after long-term treatment with rapamycin.^11,30^ Consistent with previous studies,^12–14^ short-term inhibition of the mTORC1 signaling pathway during reperfusion reduced infarct size and improved cardiac function after I/R surgery at 2 weeks (Online Figure 6).

**Figure 5.**
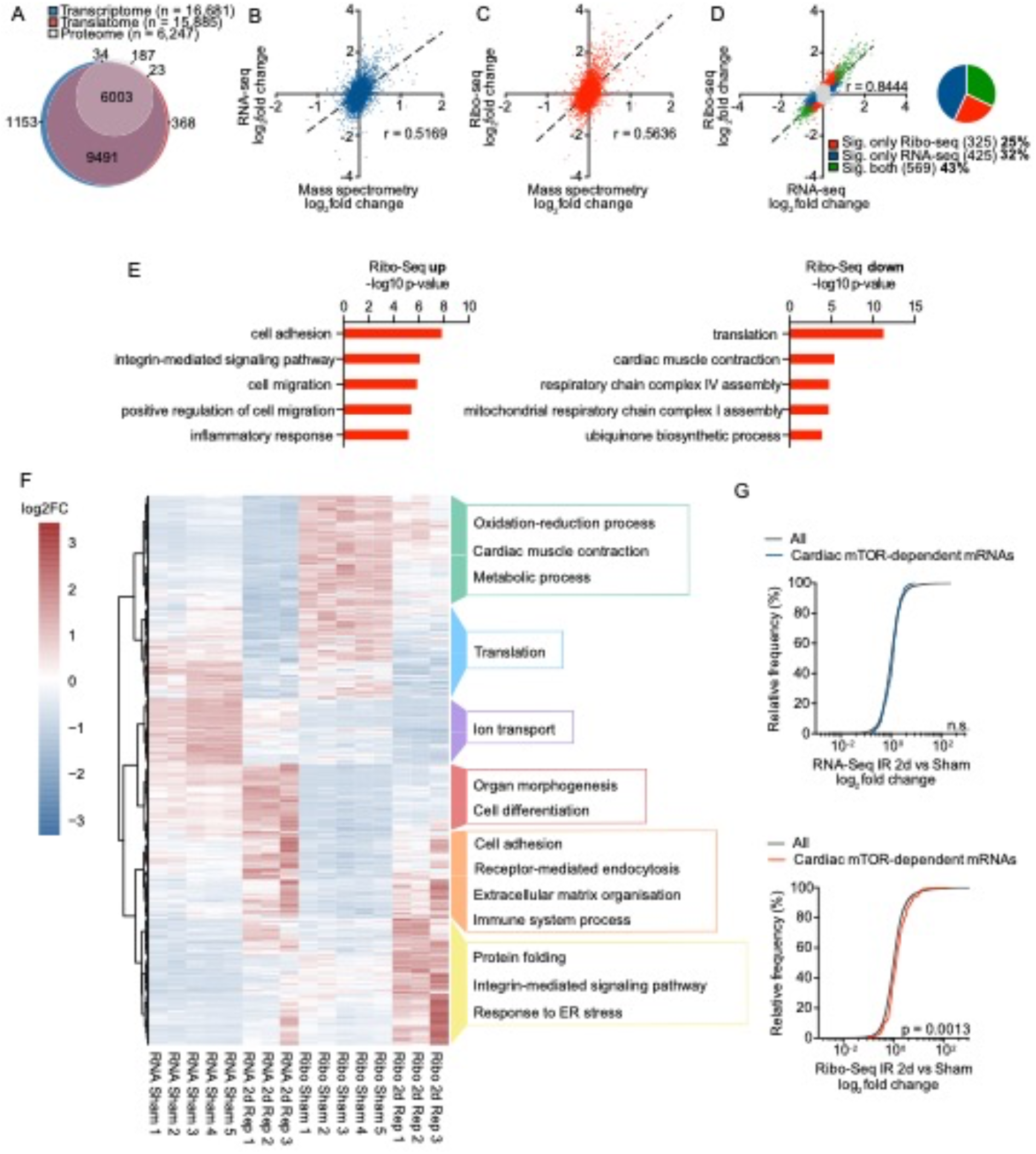
Cell-type-specific Ribo-seq identifies the translational response of cardiomyocytes to reperfusion. **A**, Venn diagram of gene products detected in the transcriptome, translatome, and proteome 2 days after I/R surgery in this study. **B** and **C**, Gene-based scatterplot showing the correlation between RNA-seq (blue, **B**) and Ribo-seq (red, **C**) expression levels and protein abundance by mass spectrometry. Correlation coefficients are Pearson r values. **D**, Scatter plot of Ribo-seq vs. RNA-seq in sham- and I/R-operated mice 2 days after surgery. Transcripts were considered significant when false discovery rate <0.05. Gray dots indicate no significant change. Significant change at translational level is shown in red, at transcriptional level in blue, and regulation at both translational and transcriptional levels in green. N ≥ 3 for each time point. **E**, Enrichment of GO terms for translationally up- and downregulated transcripts in cardiomyocytes 2d after reperfusion. The five most significant GO terms per group were displayed. **F**, Unbiased clustering analysis of RNA-seq and Ribo-Seq of differently expressed genes 2d after I/R surgery. Different colors indicate different clusters. Enriched GO terms containing more than 5 significantly regulated genes/transcripts are shown on the right for each cluster. **G**, Cumulative fraction of all detected transcripts and detected cardiac mTOR-dependent mRNAs relative to their fold change of RNA-seq (top) or Ribo-seq (below). Cardiac mTOR-dependent mRNAs were defined as genes with a heart-specific TOP score ≥ 2 from Philippe et al^50^ that were expressed in our RNA-seq and Ribo-seq datasets. RNA-seq Sham n = 5, RNA-seq Rep n = 3, Ribo-seq Sham n = 5, Ribo-seq Rep n = 3, mass spectrometry Sham = 2, mass spectrometry Rep n = 3. Rep (reperfusion). Detailed information on statistical analysis can be found in the supplementary methods.

**Figure 6.**
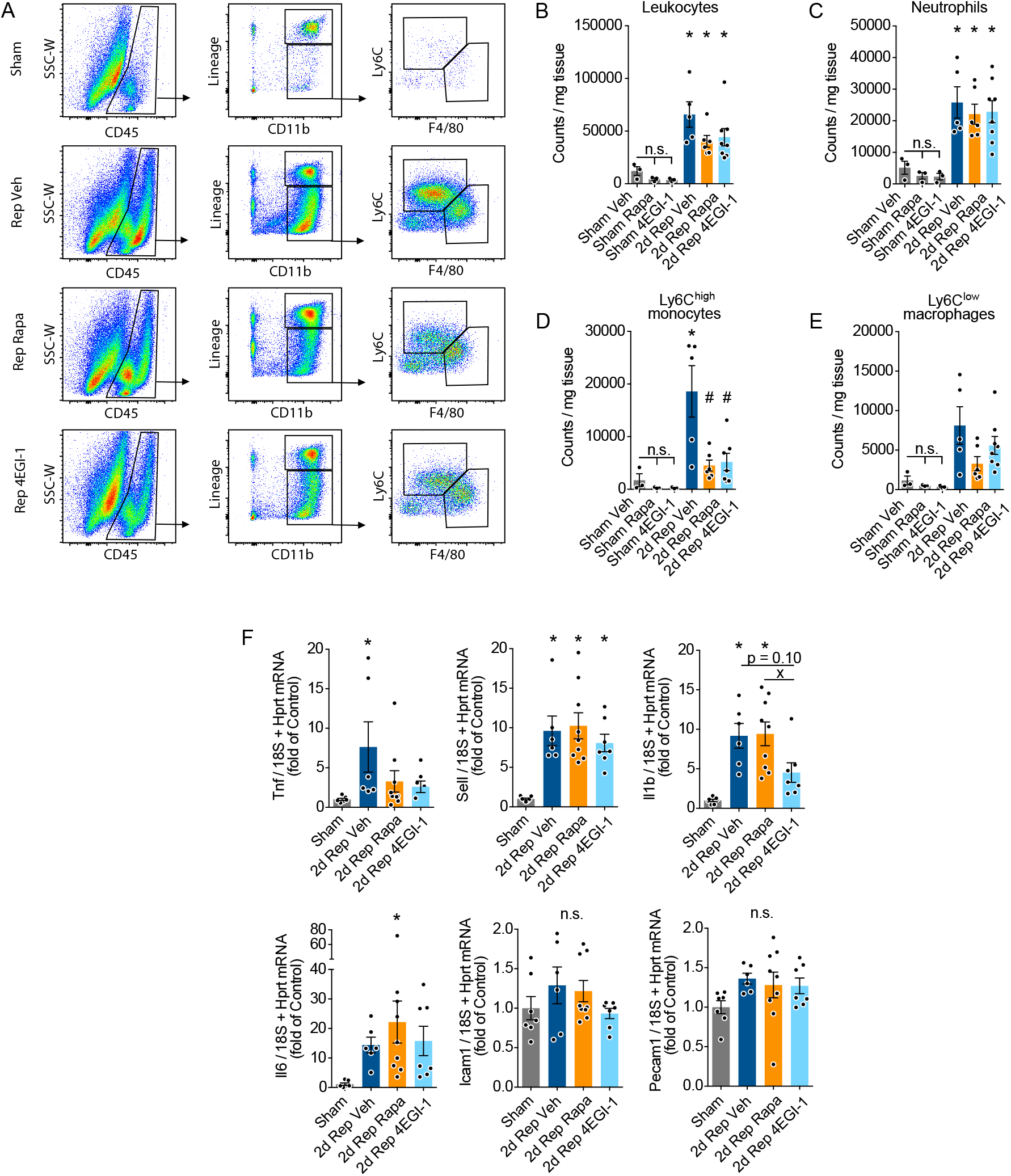
Short-term inhibition of mTORC1-4EBP1-eIF4F dependent translation during reperfusion attenuates proinflammatory Ly6C^high^ monocyte infiltration. **A**, Gating strategy and representative flow cytometric plots of mouse hearts 2 days after sham or I/R surgery treated with 2 and 6 mg/kg rapamycin and 50 mg/kg 4EGI-1. **B** to **E**, Flow-cytometry based enumeration of leucocytes (**B**), neutrophiles (**C**). Ly6C^high^ monocytes (**D**) and Ly6C^lo^/F40^+^ macrophages (**E**) per mg heart tissue 2 days after sham or I/R surgery in animals treated with vehicle, 2 and 6 mg/kg rapamycin, or 50 mg/kg 4EGI-1, n = 3-8. **F**, mRNA levels of selected inflammatory genes 2 days after sham or I/R surgery in animals treated with vehicle, 2 and 6 mg/kg rapamycin, or 50 mg/kg 4EGI-1 measured by RT-qPCR, n = 6-9. Rep (reperfusion), Veh (vehicle). * indicates p<0.05 from sham. # indicates p<0.05 from 2d Rep Veh. x indicates p<0.05 from 2d Rep Rapa. For statistical analysis one-way ANOVA with Turkey post-hoc analysis was used for **B** to **F**. p < 0.05 was defined as significant difference. Error bars show standard error of the mean (SEM).

Despite of controlling translation, mTORC1 is involved in a variety of important cellular processes, such as cellular metabolism and autophagy, which are thought regulate cardiac function in response to reperfusion.^29,31,32^ The use of 4EGI-1 allows to specifically inhibit eIF4F-dependent translation without targeting other pathways of the mTORC1 pathway, potentially circumventing side effects mediated by complete mTORC1 inhibition. Similar to our results obtained in vitro, 4EGI-1 had a low therapeutic margin in vivo. Dosing experiments revealed that 50mg/kg 4EGI-1 i.p. was the only tested dosage that significantly reduced cardiac translation without causing toxic side effects in vivo (Figure 4A and 4B). Upstream signaling of mTORC1 remained unaffected as indicated by unaltered levels of phosphorylated ribosomal protein S6 (Figure 4C and Online Figure 5). 4EGI-1 treatment of mice did not alter the amount of TUNEL^+^ cells in the peri-infarct region 2 days after I/R surgery (Figure 4D and 4E). Similar, the treatment did not affect plasma TnT levels in both female and male mice 24h after I/R surgery, indicating that 4EGI-1 does not protect from acute reperfusion injury (Figure 4F). In addition, 4EGI-1 did not improve first-day- or two-week survival (Figure 4G and 4H) or had any significant effect on heart weight to body weight ratio (HW/BW) (Figure 4I). However, similar to pharmacological mTORC1 inhibition, 4EGI-1 improved ejection fraction (Figure 4J) and reduced infarct size at 2 weeks after I/R surgery (Figure 4K). While there was a trend for reduced expression of Nppa, Nppb, Mhy7 and Col1a1, this did not reach significance (Figure 4L). No major differences were observed between female and male mice. Taken together, short-term inhibition of the mTORC1-4EBP1-eIF4F axis during reperfusion improves long-term myocardial function and reduces infarct size after cardiac I/R.

### Assessment of the cardiomyocyte-specific translatome in vivo reveals extensive translational regulation of the myocardial immune response after reperfusion

To investigate the mechanisms by which selective inhibition of translation improves long-term cardiac function after I/R surgery, we used Ribo-Seq to determine the translatome of cardiomyocytes in vivo. For this, we used our recently described approach to determine cell-type-specific translation in the heart.^6^ The translational response of mice was examined 2 days after reperfusion, the timepoint at which we observed mTORC1 activation and increased rates of translation. A comprehensive quality control of the sequencing reads was performed prior to translatome analysis (Online Figure 7). The merged translatome from all mice covered the majority of all transcribed mRNAs and almost the complete proteome of cardiomyocytes (Figure 5A). Similar to previous studies,^5,6,33^ Ribo-Seq had a higher predictive value for final protein levels after reperfusion than RNA-Seq (Pearson’s correlation coefficient [r] = 0.5636 versus 0.5169) (Figure 5B and 5C). Of all genes significantly regulated after reperfusion, approximately 25% were controlled exclusively by translation, 32% exclusively by transcription and 43% by both translation and transcription, indicating that translation regulates approximately 68% of all affected genes after reperfusion (Figure 5D).

**Figure 7.**
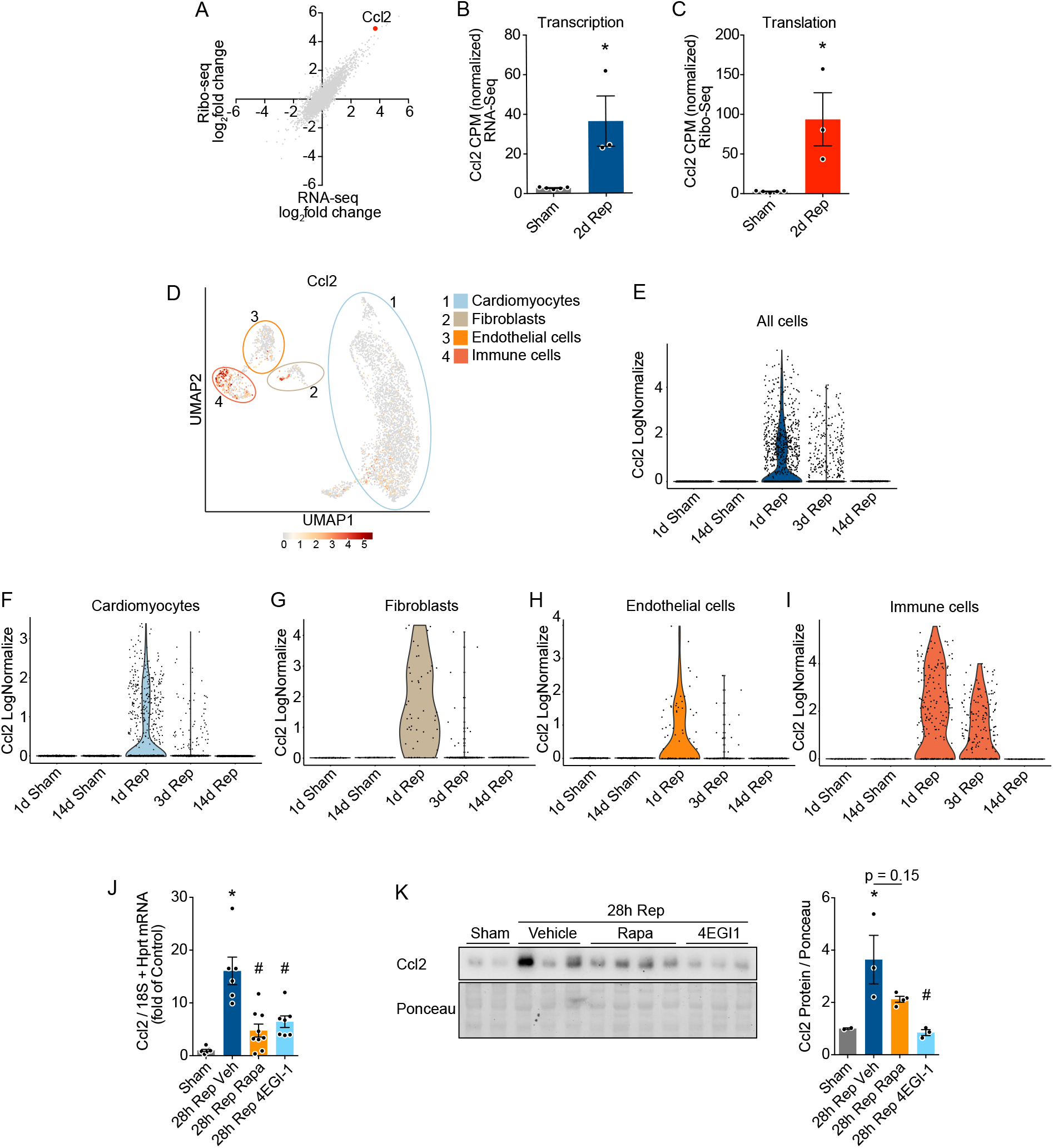
Regulation of the myocardial inflammatory response by eIF4F-dependent expression of the chemokine Ccl2. **A**, Scatter plot of Ribo-seq vs. RNA-seq in sham- and I/R-operated mice 2 days after surgery highlighting Ccl2 expression (red). **B** and **C**, RNA-seq and Ribo-seq expression data of Ccl2 in sham and I/R operated mice (CPM - count per million). **D**, Uniform manifold approximation and projection (UMAP) map indicating the expression of Ccl2 in different identified cardiac cell populations. Data are shown as normalized transcript counts on a color-coded linear scale. Different cell populations are highlighted by colored circles. **E** to **I**, Violin plots showing the mRNA expression of Ccl2 in all cells (**E**), cardiomyocytes (**F**), fibroblasts (**G**), endothelial cells (**H**) and immune cells (**I**) in sham mice or in mice after increasing timepoints after reperfusion. According to Seurat LogNormalize, gene expression measurements for each cell are normalized to total expression, multiplied by a scaling factor of 10000, and log-transformed. **J** and **K**, Ccl2 mRNA levels, measured by RT-qPCR (**J**) and Ccl2 immunoblot (**K**) of left ventricular lysates 28 hours after sham or I/R surgery in animals treated with vehicle, 2 and 6 mg/kg rapamycin, or 50 mg/kg 4EGI-1. Rep (reperfusion), Veh (vehicle). * indicates p<0.05 from sham. # indicates p<0.05 from 28h Rep Veh. Figures 7D to 7I were generated from publicly available mouse single-cell transcriptomics data published by Molenaar et al^38^. For statistical analysis one-way ANOVA with Turkey post-hoc analysis was used for **J** and **K**. An unpaired two tailed t-test was used for **B** and **C**. p < 0.05 was defined as significant difference. Error bars show standard error of the mean (SEM). Detailed information on the statistical analysis of scRNA-seq data can be found in the supplementary methods.

Cluster analysis of significantly changed genes from Ribo-Seq data revealed enrichment of cell adhesion and immune system process in transcripts upregulated after reperfusion (Figure 5E and 5F). Interestingly, mRNAs with an mTOR-dependent TOP- or TOP-like motif in the 5‵ UTR were significantly upregulated by translation, but not by transcription (Figure 5G), suggesting that the activation of the mTORC1 pathway during reperfusion results in increased translation of selected mTORC1-dependent mRNAs.

### Pharmacological inhibition of the mTORC1-4EBP1-eIF4F axis attenuates cardiac monocyte infiltration

The major signaling pathways regulated by translation in cardiomyocytes early after I/R appeared to be involved in cell infiltration and inflammation. To further characterize the myocardial inflammatory response after inhibition of the mTORC1-4EBP1-eIF4F axis during reperfusion, we examined leucocyte infiltration of hearts from mice treated with vehicle, rapamycin or 4EGI-1 48 hours after sham or I/R surgery by flow cytometry (Figure 6A). As expected, leucocytes infiltrated the heart after I/R surgery (Figure 6B). We observed a trend toward decreased overall leucocyte infiltration into the myocardium after rapamycin or 4EGI-1 treatment both at baseline and in response to I/R surgery, which did not reach significance (Figure 6B). While we did not observe any changes in neutrophil infiltration (Figure 6C), flow-cytometry based immune cell enumeration revealed severely reduced accumulation monocytes into the myocardium in response to I/R surgery after both rapamycin and 4EGI-1 treatment (Figure 6A, 6D and 6E). These changes could not be explained by blood leucocyte levels (Online Figure 8) or by differences in bone marrow proliferative cells (Online Figure 9), indicating a heart-specific response that is independent of systemic leucopoiesis.

Importantly, subpopulation gating of cardiac myeloid cells showed that this primarily affected infiltration of inflammatory Ly6C^hi^ monocytes (Figure 6D), with a lesser, non-significant effect on reparative Ly6C^lo^/F40^+^ macrophages (Figure 6E), resulting in an anti-inflammatory and pro-reparative monocyte/macrophage response to I/R. This was accompanied by blunted myocardial expression of tumor necrosis factor alpha (Tnf) after rapamycin and 4EGI-1 treatment as well as interleukin 1 beta (Il1b) exclusively after 4EGI-1 treatment (Figure 6H). The expression of selectin L (Sell), Icam1 and Pecam1 remained unaffected by both interventions (Figure 6H).

### Regulation of myocardial inflammation by mTORC1-dependent translational regulation of the monocyte-attracting chemokine Ccl2

The chemokine Ccl2, also known as monocyte chemoattractant protein-1 (Mcp-1), attracts myeloid cells to sites of injury to mediate a local inflammatory response and tissue repair.^34^ Elevated Ccl2 levels are associated with an increased risk of myocardial infarction, sudden cardiac death, coronary angioplasty, and stent restenosis.^35^ While Ccl2 was filtered from transcriptionally and translationally regulated genes after I/R surgery due to its extremely low abundance under baseline conditions (<10 counts per million reads of normalized read counts in at least one replicate), a detailed analysis that included all reads revealed that Ccl2 was among the transcripts with the highest relative increase in detected RNA-seq and Ribo-seq read abundance (Figure 7A to 7C). Previously, Ccl2 was described to be expressed primarily in non-cardiomyocytes, such as endothelial cells, fibroblasts and immune cells.^34^ However, the detection of high read-counts of Ccl2 in our cardiomyocyte-specific Ribo-seq data set indicated that cardiomyocytes express Ccl2 in response to I/R. Indeed, recent studies reported the expression and release of Ccl2 from cardiomyocytes in response to I/R by inflammatory mediators.^36,37^

To further investigate the expression of Ccl2 in cardiac cells in response to I/R in vivo, we analyzed an I/R time course of single-cell transcriptomics in mice recently published by Molenaar et al^38^ (Online Figure 10). We have made our analysis publicly available using the shinycell^39^ interactive web application accessible at https://shiny.jakobilab.org/Hofmann_et_al_2022/. Our analysis revealed that Ccl2 is expressed in several cardiac cell types, including cardiomyocytes, fibroblasts, endothelial cells, and immune cells as an early response to reperfusion injury (Figure 7D and 7E). In cardiomyocytes, Ccl2 expression was not detectable at baseline, but a subset of cardiomyocytes began to express Ccl2 in response to I/R (Figure 7F). Ccl2 expression in cardiomyocytes occurred mainly 1 to 3 days after reperfusion and decreased after 14 days, coinciding with monocyte infiltration to the heart (Figure 7F).^40^ A similar pattern was observed for other cardiac cell types, including fibroblasts, endothelial cells and immune cells (Figure 7G to 7I). Importantly, the expression of Ccl2 was greatly attenuated after I/R in response to inhibition of the mTORC1-4EBP1-eIF4F axis by rapamycin or 4EGI-1 (Figure 7J), which we confirmed on the protein level (Figure 7K). In summary, inhibition of eIF4F dependent translation reduced inflammatory monocyte infiltration at least in part by inhibiting cardiac Ccl2 expression.

We conclude that a transient pharmacological intervention that inhibits translation by targeting the mTORC1-4EBP1-eIF4F axis at the onset of reperfusion improves long-term cardiac function at least partly by tempering the reperfusion-driven inflammatory monocyte response via lowered expression of Ccl2.

## Discussion

Ischemia critically limits oxygen and nutrient availability, resulting in cell death and loss of viable myocardium. Rapid reperfusion of the ischemic myocardium is the most effective treatment for acute myocardial infarction.^2^ However, reperfusion itself contributes significantly to myocardial damage known as reperfusion injury^3^. Currently, no intervention targeting reperfusion injury has been successfully translated to the clinic, highlighting the critical need for a detailed understanding of the pathophysiology of the myocardial response to reperfusion.^3^ Here, we describe a maladaptive translational response of border zone cells, including cardiomyocytes, mediated by the mTORC1-4EBP1-eIF4F pathway that promotes myocardial immune cell infiltration and inflammation. This pathway can be specifically targeted by a pharmacological intervention immediately before and after reperfusion to improve long-term cardiac function after acute myocardial infarction.

### Dynamic regulation of translation and cardiac function in response to I/R via the mTORC1-4EBP1-eIF4F axis

Translation is a highly energy-dependent process and thus heavily regulated by cellular availability of nutrients, ATP and oxygen.^41^ While translation has been extensively studied in the context of ischemia, relatively little is known about the regulation of translation in response to cardiac reperfusion. Ischemic preconditioning, that is, brief episodes of ischemia followed by reperfusion that increases the resistance of the myocardium to subsequent ischemic insults, has been shown to be associated with a rapid increase in protein synthesis in the ischemic-reperfused region.^42^ Furthermore, Zhang et al^43^ recently described the transient suppression of protein synthesis in H9c2 cardiomyoblasts and mice immediately after reperfusion, which we reproduced here in NRCMs. This observation could possibly be explained by additional injury immediately after cellular reperfusion.

We identified an additional upregulation of myocardial translation 48h after reperfusion in the border zone. We further demonstrated that plasma TnT levels were positively correlated to the amount of puromycin incorporation 2 days after reperfusion, suggesting that the intensity of the translational response is directly linked on the amount of injured myocardium. Using a combination of pharmacological and genetic methods, we showed that regulation of cardiomyocyte translation in response to ischemia and reperfusion is a direct consequence of changes in the activity of the mTORC1-4EBP1-eIF4F pathway. Furthermore, we confirmed that eIF4E^S209^ phosphorylation does not affect the total amount of translation or I/R injury. This is consistent with the conclusion of several previous reports that neither MNK1/2 kinases nor eIF4E^S209^ phosphorylation are essential for regulating overall translation rates.^44–46^ Similar to the mTORC1-4EBP1-eIF4F pathway, the integrated stress response was recently shown to regulate translation initiation via the PERK-eIF2α axis in response to cardiac reperfusion.^43^ Thus, both the mTORC1-4EBP1-eIF4F and PERK-eIF2α pathways, as well as possibly other signaling pathways, appear to jointly control translation initiation after reperfusion. The activity of these different signaling pathways could specifically affect the translational efficiency of a different subset of transcripts after reperfusion, which could be further investigated in future studies.

### Translational regulation of myocardial inflammation in response to reperfusion

To gain further insight into the functional consequence of activation of cap-dependent translation in the border zone in response to myocardial reperfusion, we treated mice with 4EGI-1, a competitive eIF4E/eIF4G interaction inhibitor that increases binding of 4EBP1 to eIF4E. 4EGI-1 efficiently inhibited upregulation of translation after reperfusion, reduced infarct size and improved ejection fraction 2 weeks after myocardial infarction. These results highlight the importance of translational regulation for the protective effects mediated by pharmacological mTORC1 inhibition, as recently shown for other cardiac disease.^9^ Caution should be taken regarding the potential toxic off-target effects of 4EGI-1, which we believe severely limit its potential clinical application in its current state. However, the notion that transient inhibition of translation may protect against cardiac I/R injury is consistent with previous observations that the translational inhibitor cycloheximide protects against cell death in an ex vivo model of I/R^47^ and with recent findings by Zhang et al, showing that activation of the integrated stress response protects the heart from reperfusion injury in vivo.^43^

The upregulation of myocardial translation at the onset of reperfusion appeared to be mediated predominantly by border zone cells, including cardiomyocytes. We therefore determined the cardiomyocyte-specific translational response of mice subjected to I/R reperfusion in vivo, which revealed extensive regulation of the inflammatory response by cardiomyocytes at the level of translation. Inhibition of cardiac eIF4F-dependent translation specifically inhibited myocardial accumulation of pro-inflammatory monocytes after I/R injury. We identified the monocyte-attracting chemokine Ccl2 as a target regulated by the mTORC1-4EBP1-eIF4F pathway that was expressed by a subset of cardiomyocytes exclusively in response to reperfusion, as has been recently described for isolated cardiomyocytes.^36,37^ Inhibition of the CCR2-CCL2 axis was previously shown to exhibit beneficial effects in murine models of myocardial infarction.^48^ In addition, mTORC1 was previously described to regulate

Ccl2 secretion in immune cells.^49^ Our study primarily examined the translational response of cardiomyocytes to I/R. While cardiomyocytes were shown to regulate a maladaptive protein synthesis response after reperfusion, other cell types are likely to be affected by our systemic pharmacological intervention. Our single-cell transcriptomics analysis revealed the contribution of several cell types to cardiac Ccl2 expression, which included fibroblasts, endothelial cells, and immune cells. In addition, mTORC1 inhibitors such as rapamycin are widely used immunosuppressive drugs due to their broad spectrum of inhibitory effects on immune cells.^49^ Therefore, it seems likely that the adaptive effects of mTORC1 and eIF4F inhibitors in response to I/R are mediated by a spectrum of anti-inflammatory actions across different cell types. Nevertheless, pharmacological inhibition of the mTORC1-4EBP1-eIF4F pathway mediated its protective effect, at least in part, via control of Ccl2 expression in cardiomyocytes and subsequent regulation of inflammatory monocyte infiltration into the heart.

In conclusion, we identified an unexpected maladaptive translational response of the myocardial border zone mediated by the mTORC1-4EBP1-eIF4F axis that controls cardiac inflammation. A short-term pharmacological intervention that inhibits eIF4F-dependent translation after reperfusion improves long-term cardiac function, caused at least in part by regulating inflammatory monocyte infiltration to the heart via Ccl2.

## Acknowledgments

C.H. and M.V. conceived and designed the study. C.H., A.S., O.M.S., F.S.Y., F.S., I.S.M., E.M., C.Sa., L.J., V.K. and C.St. performed the experiments. C.H., A.S., O.M.S., F.S.Y., M.R., F.S. and T.J. analyzed the data. N.F., H.A.K. and M.V. acquired funding. C.H. and M.V. supervised the study. C.H. and M.V. visualized the data; C.H. contributed to writing—original draft; C.H., H.A.K., N.F., F.L. and M.V. contributed to writing—review and editing. All authors approved the final version of the article. The authors would like to acknowledge the expertise and support of E. Boileau for assistance with sequencing data analysis.

## Sources of Funding

C.H. acknowledges the German Cardiac Society (DGK). E.M. acknowledges the Heidelberg Biosciences International Graduate School (HBIGS). H.A.K., N.F., F.L. and M.V acknowledge the DZHK (German Centre for Cardiovascular Research) Partner Site Heidelberg/Mannheim. M.V. acknowledges the DFG (German Research Foundation, DFG VO 1659 2/1, DFG VO 1659 2/2, DFG VO 1659 4/1, DFG VO 1659 6/1) and the Boehringer Ingelheim Foundation (Plus 3 Program).

## Disclosures

None.

## Notes

### Competing Interest Statement

The authors have declared no competing interest.

### Summary of Updates

-

