## Supplemental Material for "Transient inhibition of translation improves long-term cardiac function after ischemia/reperfusion by attenuating the inflammatory response"

[5] Lead contact

#### Expanded Materials and Methods

##### Cultured Cardiomyocytes

Isolation of neonatal rat ventricular cardiomyocytes (NRCMs) were isolated as described before.[1] Cells were isolated from one to two-day old WISTA rats (cat# 13792727, WISTAR HanRj:WI, Janvier Labs) via enzymatic digestion and purified by Percoll density gradient centrifugation. Cardiac myocytes were then plated at a density of 0.5 X $10^6$ cells per well on 34.8 mm plastic plates that had been pre-treated with 5 µg/ml fibronectin (cat# F1141, Sigma-Aldrich) in serum free DMEM/F12 medium (cat# 11330032, Thermo Fisher Scientific) for one hour and then cultured in DMEM/F12 1:1, containing 10% fetal bovine serum, 100 units/mL of penicillin, 100µg/mL streptomycin and 292 µg/ml glutamine (cat# 10378016, Thermo Fisher Scientific). After 24h, media was changed once to remove dead and non-adherent cells. After additional 24h cells were subjected to the respective treatments.

##### Simulated Ischemia/Reperfusion in Vitro

Cells were cultured as described above. 48h-72h after isolation cells were washed twice with PBS and medium was changed to low-nutrient DMEM without D-glucose, sodium pyruvate, HEPES, L-glutamine and phenol red (cat# A1443001, Thermo Fisher Scientific), supplemented with 0.5% dialyzed fetal bovine serum (cat# A3382001, Thermo Fisher Scientific), 100 units/ml penicillin, 100µg/ml streptomycin, and 292 µg/ml glutamine. Cells were then incubated in a hypoxia incubator at 37°C, 5.0% $CO_2$ and 0.2% $O_2$ for one to 24 hours. For subsequent simulated reperfusion, medium was changed to DMEM/F-12 supplemented with 10% fetal bovine serum and 100 units/mL of penicillin, 100µg/mL streptomycin and 292 µg/ml glutamine, followed by incubation in an incubator at 37°C, 5.0% CO2 and 21% $O_2$. Control cells were cultured in DMEM/F-12 supplemented with 10% fetal bovine serum and 100 units/mL of penicillin, 100µg/mL streptomycin and 292 µg/ml glutamine at 37°C, 5.0% CO2 and 21% $O_2$ for as long as the longest ischemia/reperfusion timepoint.

##### Laboratory Animals

Data associated with all animal studies reported in this article has been reviewed and approved by the institutional animal care and use committee. All experiments were performed in 10- to 12-week-old male or female C57BL/6N mice. All animals were fed *ad libitum* and were housed at Heidelberg University in a temperature- and humidity-controlled facility with a 12-h light-dark cycle. Cardiomyocyte-specific Ribo-tag mice were previously described.[2] Briefly, Ribo-tag mice (JAX ID 011029) were bred to αMHC-Cre mice to obtain Rpl22-HA-expressing homozygous mice in cardiomyocytes. Rapamycin was diluted in 100 µl PBS or 15µl DMSO and 200µl corn oil. 4EGI-1 was diluted in 15µl DMSO and 200µl corn oil. Drugs were administered by i.p. injection. Dosing studies were performed with 6 mg/kg rapamycin or 10 to 100 mg/kg 4EGI-1. In I/R-operated mice receiving drug treatment, mice received a first dose of 6 mg/kg rapamycin or 50 mg/kg 4EGI-1 30 minutes before reperfusion and a second dose of 2 mg/kg rapamycin or 50 mg/kg 4EGI-1 24 hours after surgery. Control mice in these experiments received equal amounts of vehicle at the same time points.

**Ischemia/Reperfusion and Myocardial Infarction Surgery in Vivo**

Animals were randomly assigned to each experimental group. The mice received a s.c. injection of 0.1 mg/kg buprenorphine and were anesthetized after 30 min with a 5% isoflurane/$O_2$ mixture using an anesthetic vaporizer (Harvard Apparatus). Then, mice were fixed on a warming plate heated to 37°C and intubated. The intubation catheter was connected to a ventilator (MiniVent type 845, Harvard Apparatus), and the respiratory volume was set to approximately 260µl and the respiratory rate to approximately 180/min. A grease cream was applied to the eyes to protect them from dehydration. After successful intubation, the isoflurane was reduced to 1.5%. A depilatory cream was used to first depilate the thorax of the mice. Each mouse received 250µl of sterile NaCl solution subcutaneously before surgery. A skin incision of approximately 1.5cm was made to the left of the xiphoid in the direction of the left axilla. Subsequently, the fasciae of the pectoralis muscles were exposed. The pectoralis muscles were pierced with a 6-0 Prolene suture on their lateral ends and fixed next to the mouse on the heating pad, exposing the ribs and intercostal muscles. Then, the intercostal muscle of the third intercostal space was pierced with the tip of forceps and incised using microsurgical scissors. Care was taken not to injure the internal thoracic artery and the lung. After the incision, another 6-0 Prolene suture was pierced around the third rib and fixed cranially of the mouse to spread the ribs. Now the exposed pericardium was carefully dissected apart, and the thymus was pushed aside. Next, an 8-0 Prolene suture was pierced under the left anterior descending artery approximately 1mm caudal to the left atrium and perpendicular to the course of the vessel for a length of approximately 3 mm. Ischemia was induced by knotting a suture over a PE-10 tube which was placed over the vessel, thereby compressing the lumen of the artery. Ischemia was maintained for 60 minutes. During the period of ischemia, isoflurane was reduced to 0.75% - 1% and the open thorax was covered with a warmed swab moistened with NaCl. If drugs or vehicles were to be administered, the first dose was injected i.p. 30 minutes before reperfusion. For reperfusion, isoflurane was increased to 1.5%, the cover was removed, and the PE-10 tube was pulled out. Then the 8-0 Prolene suture was cut open and removed. The pericardium was positioned back on the heart and the chest was closed with one to two sutures. The skin was closed with three to four simple interrupted stitches. The surgical wound was cleaned with saline. Isoflurane was now stopped, and the mouse continued to be ventilated with 100% oxygen until it awoke. Once it regained its own respiratory activity, it was extubated and placed in a warmed chamber flooded with 100% oxygen until it was fully awake and moving again. Post-operatively, the mouse received 0.1mg/kg buprenorphine s.c. every 6h for 72h. Myocardial infarction surgery was performed by permanently ligating the left anterior descending artery according to the procedure described above. Sham surgeries were performed following the same procedures except that the LAD was not ligated.

**Plasma Troponin T and CK Measurement**

Twenty-four hours after surgery, the mouse is brought to brief anesthesia with moderate respiratory slowing with isoflurane to allow retrobulbar blood sampling. Heparin-coated capillaries were used for retrobulbar blood sampling. Blood samples were kept on ice and centrifuged for 10 minutes at 1000rcf. 10µl of plasma was taken from the supernatant and diluted 1:30 in PBS. Troponin T and CK analysis was performed at the central laboratory of Heidelberg University Hospital.

**Echocardiography**

Echocardiography was carried out on anesthetized mice using a Visualsonics Vevo 2100 high-resolution echocardiograph. Anesthesia was administered via a facial mask and maintained by a minimum dose of isoflurane (1.0–2.0%). Echocardiography was performed at a heart rate of 450-550 bpm.

**Preparation of Tissue Lysates**

Mice were sacrificed, and left ventricles were rapidly excised, washed in PBS and snap frozen in liquid nitrogen. Tissue used for Ribo-Seq and RNA-Seq analysis was washed in PBS containing 100 µg/ml cycloheximide. Left ventricles were homogenized using a tissue homogenizer in 5 volumes of ice-cold polysome buffer containing 20mM Tris pH 7.4, 10mM MgCl, 200mM KCl and 1% Trition X-100. Tissue used for Ribo-Seq and RNA-Seq analysis was homogenized in polysome buffer containing 20 mM Tris pH 7.4, 10 mM MgCl, 200 mM KCl, 2 mM DTT, 1% Triton X-100, 1U DNAse/µl and 100 µg/ml CHX. For library construction lysates were processed as previously described.[2] For protein analysis via immunoblotting, initial lysates were further diluted with 9 volumes of RIPA buffer containing 20mM Tris-HCl (pH 7.4), 150mM NaCl, 1% Triton X-100, 0.1% SDS, 0,5% Sodium deoxycholate, protease inhibitor cOmplete ULTRA (cat# 05892791001, Roche) and phosphatase inhibitor PhosSTOP (cat# 04906837001, Sigma-Aldrich). RNA was isolated from tissue lysates using TRIzol (cat# 15596026, Invitrogen).

**Puromycin Incorporation Assay**

Changes in the amount of protein synthesis of differentially treated cells were followed by puromycin incorporation. Puromycin is a structural analogue of aminoacyl-transfer RNA and can be incorporated into elongating peptide chains which causes premature release of truncated puromycin bound peptides from the ribosome. At very low concentrations, that do not inhibit the overall rate of translation, the rate at which puromycin peptides are formed reflects the overall rate of protein synthesis. In vitro 0.5 µg/ml was added to the culture medium 30 minutes before harvesting the cells. Thereafter, cells were washed once with ice-cold PBS and then harvested as described above. To assess translation rates in vivo, mice were i.p. injected with 50mg/kg puromycin or O-propargyl-puromycin 30 minutes before the animals were sacrificed. O-propargyl-puromycin was detected with the Click-iT Plus OPP Alexa Fluor™ 594 Protein Synthesis Assay Kit (Invitrogen) according to manufacturer's instructions.

**m7GTP Immunoprecipitation**

To enrich mRNA cap associated proteins from cell lysates, agarose beads coupled to 7-Methylguanosine-5'-triphosphate (m7GTP) were used, which mimics the mRNA 5´ cap structure. Cells were washed once with ice-cold PBS and harvested in 600µl NP-40 lysis buffer supplemented with 1x PhosSTOP and 1x protease inhibitor cOmplete ULTRA. Maximum lysate volume of the sample with the lowest protein concentration was taken as input for the following pull-down, to which all other samples were normalized (around 2000 - 3500µg, ~ 500µl lysate). The corresponding input volumes were filled up to 700µl with NP-40 lysis buffer to ensure sufficient rotation of the beads during incubation. 40µl of the lysates were kept as an input control for western blotting to estimate pull-down efficiency. For the following steps, low retention tubes were used. For each sample 50µl of m7GTP agarose beads were used. To equilibrate the beads, they were washed three times with 1 ml cold NP-40 lysis buffer and centrifuged at 500 G and 4°C for 3 minutes. After the last washing step, the beads were resuspended 1:2 in cold NP-40 lysis buffer and 150µl of the diluted beads were added to the prepared samples, then they were rotated (10 rpm) over night at 4°C. The next day, the samples were centrifuged at 500 G and 4°C for 3 minutes, the supernatant was discarded, and the beads were washed three times with 1ml cold NP-40 lysis buffer. To separate bound proteins from the beads, 40µl 1x Laemmli Sample Buffer with 10% β-Mercaptoethanol were added to each sample and the samples were cooked at 95°C for 5 minutes. After centrifugation at 500 G for 3 minutes, the supernatant was collected, and a western blot was performed. For each sample, maximal volume of the respective supernatant as well as 30µg of the input control were loaded.

**Immunoblotting**

Cultured cells were lysed in RIPA Buffer consisting of 50mM Tris pH 7.5, 150mM NaCl, 1% Triton X-100 and 1% SDS, which was supplemented with protease inhibitor cOmplete ULTRA (cat# 05892791001, Roche) phosphatase inhibitor PhosSTOP (cat# 04906837001, Roche). Tissue lysates were prepared as described above. Lysates were cleared by centrifugation at 4 °C for 10 minutes at 20.000 rcf. Lysate protein concentration was determined using the DC Protein Assay Kit II (cat# 5000112, Bio-Rad) according to the manufacturer's instructions. Equivalent amounts of protein, usually 20-30 µg, were brought up to similar volume, mixed with Laemmli Sample Buffer (Bio-Rad; 161-0747) and 2-Mercapoethanol (cat# M6250, Sigma-Aldrich) and boiled at 95°C for 5 minutes. Samples were separated on SDS-PAGE gels and transferred to Immobilon-P transfer membranes (cat# IPVH00010, Merck Millipore). The following antibodies were used to probe the membranes: eIF4G (cat# 2498, Cell Signaling Technology, 1:5000), eIF4A (cat# C32B4, 2013, Cell Signaling Technology, 1:1000), Phospho-eIF4E Ser209 (cat# 9741, Cell Signaling Technology, 1:1000), eIF4E (cat# 9742, Cell Signaling Technology, 1:5000), Phospho-p70 S6 Kinase Thr389 (cat#9205, Cell Signaling Technology, 1:1000), p70 S6 Kinase (cat# 49D7, 2708, Cell Signaling Technology, 1:1000), Phospho-Akt Ser473 (cat# 9271, Cell Signaling Technology, 1:1000), Phospho-Ribosomal S6 Ser235/236 (cat# D57.2.2E, 4858, Cell Signaling Technology, 1:5000), Ribosomal S6 Ser235/236 (cat# 54D2, 2317, Cell Signaling Technology, 1:1000), Phospho-4EBP1 Thr37/46 (cat# 236B4, 2855, Cell Signaling Technology, 1:5000), 4EBP1 (cat# 9452, Cell Signaling Technology, 1:5000), P-AMPK (cat# 2535, Cell Signaling Technology, 1:1000), AMPK (cat# 2532, Cell Signaling Technology, 1:1000), Phospho-ERK1/2 Thr202/Tyr204 (cat# 9101, Cell Signaling Technology, 1:5000), Phospho-p38 MAPK Thr180/Tyr182 (cat# D3F9, 4511, Cell Signaling Technology, 1:1000), Phospho-Hsp27 Ser82 (cat# D1H2F6, 9709, Cell

Signaling Technology, 1:1000), Ccl2 (cat# 66272-1-Ig, 1B9F7, proteintech, 1:1000), Puromycin (cat# MABE343, Merck Millipore; 1:10,000-50,000), GAPDH (cat# G-9, sc-365062, Santa Cruz Biotechnology; 1:20,000), β-Actin (cat# C4, sc-47778, Santa Cruz Biotechnology; 1:20,000), Ponceau solution was prepared with Ponceau BS (cat# B6008, Sigma Aldrich).

**Quantitative Real Time PCR**

Total RNA was isolated from cultured cardiomyocytes using the Quick-RNA MiniPrep Kit (cat# R1055, Zymo Research) and from tissue using the RNeasy Mini Kit (cat# 74104, Qiagen) according to the manufacturer's instructions. cDNA was generated by reverse transcription using Superscript III First-Strand Synthesis System (Invitrogen; 18080-051). Quantitative Real Time PCR was performed with Maxima SYBR Green/ROX qPCR Master Mix (Thermo Fisher cat# K0222) in a StepOnePlus RT-PCR System (Thermo Fisher). The following primers were used:

Mouse-CCL2-F: CACTCACCTGCTGCTACTCA

Mouse-CCL2-R: TTGAGCTTGGTGACAAAAACTACA

Mouse-CCR2-F: AGGAGCCATACCTGTAAATGCC

Mouse-CCR2-R: ATGCCGTGGATGAACTGAGG

Mouse-ICAM-1-F: CCCACGCTACCTCTGCTC

Mouse-ICAM-1-R: GATGGATACCTGAGCATCACC

Mouse-IL-1β-F: AGCTGGATGCTCTCATCAGG

Mouse-IL-1β-R: AGTTGACGGACCCCAAAAG

Mouse-IL6-F: GATGCTACCAAACTGGATATAATC

Mouse-IL-6-R: GGTCCTAGCCACTGGATCTGTG

Mouse-mSelectin-F: TGGTCATCTCCAGAGCCAAT

Mouse-mSelectin-R: GCAGTCCATGGTACCCAACT

Mouse-PECAM-1-F: CGGTGTTCAGCGAGGTCC

Mouse-PECAM-1-R: ACTCGACAGGATGGAAATCAC

Mouse-TNFα-F: CCATTCCTGAGTTCTGCAAAG

Mouse-TNFα-R: GCAAATATAAATAGAGGGGGGC

Mouse-VCAM1-F: TCTTACCTGTGCGCTAATGAGT

Mouse-VCAM1-R: ACTGGATCTTCAGGGAATGAG

**Immunofluorescence of Mouse Heart Sections**

Immunocytofluorescence of cardiac sections was performed as previously describe.[2] Briefly, hearts were retroperfused with PBS at 70 mmHg, arrested in diastole with 60mM KCl, fixed by perfusion for 15 minutes with 10% formalin (Sigma; HT501128), excised and fixed in formalin for 24 hours at room temperature. Fixed hearts were then dehydrated and paraffinized using a HistoCore Pearl and Arcadia H, sectioned and placed on glass slides. For immunofluorescence staining, the sections were deparaffinized and boiled in 10mM citrate buffer (pH 6.0) for 12 min. After cooling, the sections were washed 2x for 5 min with PBS and then treated with TNB blocking solution (0.5% TSA blocking reagent dissolved in TN buffer (0.1 M Tris-HCl (pH 7.5), 0.15 M NaCl)) for 60 minutes. Sections were stained with antibodies against Troponin T (cat#ab209813, Abcam, 1:300) or Phospho-S6 Ribosomal Protein (Ser235/236) (D57.2.2E) XP® Rabbit mAb (Alexa Fluor® 488 Conjugate) (cat#4803, Cell Signaling Technology), DAPI OP-Puromycin (Click-iT™ Plus OPP Alexa Fluor™ 594 Protein Synthesis Assay Kit, cat#C10457, Invitrogen, according to the manufacturer's protocol), TUNEL staining (In Situ Cell Death Detection Kit, Fluorescein, cat#11684795910, Roche, according to the manufacturer's protocol). The secondary antibody used was Cy3 AffiniPure Donkey Anti-Rabbit IgG (H+L) (cat# 711-165-152, Jackson ImmunoResearch, 1:100).

**TUNEL Cell Death Assay**

Cardiac sections were prepared as described above. The TUNEL staining was performed according to the manufacturer's protocol (In Situ Cell Death Detection Kit, Fluorescein, cat#11684795910, Roche). For each heart, 10 randomly selected images with an edge length of 488µm were acquired near the infarct area and TUNEL-positive cells were quantified using ImageJ.

**Infarct Size Quantification**

Isolated hearts were cut into five transversal sections and further processed as described above. Cardiac sections were then stained with Masson's trichrome stain. An Axio Vert. A1 microscope was used to take pictures of each heart. The percentage of the blue fibrous area to the total area of the cardiac section was defined as infarct area. ImageJ was used to determine the infarct area using the threshold area. The investigator was blinded to treatment during quantification.

**Flow cytometry – Apoptosis Assay in Vitro**

Apoptosis of isolated cardiomyocytes was induced using $H_2O_2$ (50 µM) for 4 hours. Cells were treated with the appropriate inhibitors during $H_2O_2$ treatment. Apoptosis was quantified using the Dead Cell Apoptosis Kits with Annexin V for Flow Cytometry (cat# V13242, Invitrogen) according to the manufacturer's protocol. Flow cytometry was performed using a FACS verse. Data were analyzed using FlowJo v10.

**Flow cytometry – Immune Response in Vivo**

Hearts were harvested 48 hours after I/R surgery, cleared of blood and placed in 1ml PBS on ice. To obtain a single cell solution, hearts were cut into small pieces and digested in 1ml digestion solution consisting of 1% collagenase XI, 0.5% hyaluronidase, 4.5% collagenase I, 0.3% DNase, 2% 1M HEPES and PBS at 37°C for 60 minutes. The digestion solution was filtered through a 40µm cell strainer with 40ml FACS buffer consisting of 2% FCS, 2mM EDTA and PBS and centrifuged at 4°C and 800 rcf for 5 minutes. Cells were washed twice with FACS buffer and then stained with Ly6C, F4/80, CD45, CD11b, and lineage (Ter119, CD90, B220, CD49b, NK1.1, Ly6G) for 30 minutes, each diluted 1:200 at 4°C. Afterwards, the staining was stopped using 900µl FACS buffer and the cell solution was transferred to a counting tube after re-centrifugation and resuspension in 1ml FACS buffer. Flow cytometry was performed using a FACS verse. Data were analyzed using FlowJo v10. Gating strategy was performed as previously described.[3]

**Parallel Generation of Ribo-seq and RNA-seq Libraries**

Libraries were generated as previously described.[2] For each animal, the heart was lysed in 700µl polysome buffer (20 mM Tris pH 7.4, 10 mM MgCl, 200 mM KCl, 2 mM DTT, 1% Triton X-100, 1U DNAse/µl and 100 µg/ml CHX) using a tissue homogenizer (Bullet Blender, NextAdvance). The tissue was homogenized further by passing the lysate through a 23-gauge syringe needle ten times. Homogenates were centrifuged at 4°C and 18,000xg for 10 min, and the supernatant was immediately used in the further steps. For complete lysis, the samples were kept on ice for 10 min and subsequently centrifuged at 20,000xg to precipitate cell debris. To accurately dissect translation and transcription, both Ribo-seq and RNA-seq libraries were prepared for each biological replicate from the identical lysate. Ribosome footprints were generated after immunoprecipitation of cardiomyocyte-specific polysomes with Anti-HA magnetic beads after treating the lysate with RNAse I (Ambion). Libraries were generated according to the mammalian Ribo-seq kit (Illumina). Barcodes were used to perform multiplex sequencing and create sequencing pools containing at least eight different samples and always an equal amount of both RNA and ribosome protected fragments (RPF) libraries. Sample pools were sequenced on the HiSeq 2000 platform using 50-bp sequencing chemistry.

**Sequencing Data Processing and Quality Control**

Sequencing data processing and quality control was performed as previously described.[2] Adapters removal was done with Flexbar v3.0.3[4] using standard filtering parameters (no prior trimming). Reads with more than 1 uncalled base were not included in the output: flexbar *--may-uncalled 1 --pre-trim-left 0*. Reads aligning to a custom bowtie2 v2.3.0[5] index including mouse rRNA, mtRNA, tRNA, snRNA and other ncRNA (Ensembl mus muculus release 97 – ncrna) were discarded. Remaining reads were then aligned in genomic coordinates to the mouse genome (GRCm38.p6) with STAR, v.2.5.3a[6], inserting annotations on the fly: STAR *--quantMode TranscriptomeSAM --alignIntronMin 20 --alignIntronMax 100000 --outFilterMismatchNmax 1 --outFilterIntronMotifs RemoveNoncanonicalUnannotated --outFilterMismatchNoverLmax 0.04 --sjdbOverhang 50*. Only uniquely mapping reads were kept for analysis. For Ribo-seq data, only periodic fragment lengths were kept that showed a distinctive triplet periodicity. We used the automatic Bayesian selection of read lengths and ribosome P-site offsets (BPPS) method[7] to select and shift aligned reads to properly account for P-site of the ribosome. For the RNA-seq data, reads were trimmed from the 3' end after adapter removal, such that the read length before alignment did match the maximum periodic fragment length of the corresponding Ribo-seq sample, as determined with the BPPS method. Finally, abundance estimates and read count to coding sequences were obtained using HTSeq-count[8], taking into account the strand-specific protocols. edgeR with standard parameters was used for differential gene expression analysis. We used an FDR<0.05 as cutoff.

**Gene Ontology Analysis**

GO term enrichment analysis was performed as previously described. Briefly, genes with FDR <0.05 were considered for further analysis. GOTERM_BP_DIRECT in DAVID v6.8[9,10] was used with the subset of expressed protein-coding genes as background set. Only enriched GO terms with at least five significantly changed genes were kept for further analysis and ranked by p-value. Top enriched terms were retained and visualized with a custom plotting routine showing enrichment of p-value.

**Mass Spectrometry Sample Preparation SP3 and TMT labeling, OASIS**

Reduction of disulfide bridges in cysteine containing proteins was performed with dithiothreitol (56°C, 30 min, 10 mM in 50 mM HEPES, pH 8.5). Reduced cysteines were alkylated with 2-chloroacetamide (room temperature, in the dark, 30 min, 20 mM in 50 mM HEPES, pH 8.5). Samples were prepared using the SP3 protocol[11] and trypsin (sequencing grade, Promega) was added in an enzyme to protein ratio 1:50 for overnight digestion at 37°C. Next day, peptide recovery in HEPES buffer by collecting supernatant on magnet and combining with second elution wash of beads with HEPES buffer.

Peptides were labelled with TMT10plex[12] Isobaric Label Reagent (ThermoFisher) according to the manufacturer's instructions. Samples were combined for the TMT10plex and for further sample clean up an OASIS® HLB µElution Plate (Waters) was used. Offline high pH reverse phase fractionation was carried out on an Agilent 1200 Infinity high-performance liquid chromatography system, equipped with a Gemini C18 column (3 µm, 110 Å, 100 x 1.0 mm, Phenomenex).[13]

**LC-MS/MS**

An UltiMate 3000 RSLC nano LC system (Dionex) fitted with a trapping cartridge (µ-Precolumn C18 PepMap 100, 5µm, 300 µm i.d. x 5 mm, 100 Å) and an analytical column (nanoEase™ M/Z HSS T3 column 75 µm x 250 mm C18, 1.8 µm, 100 Å, Waters). Trapping was carried out with a constant flow of 0.05% trifluoroacetic acid at 30 µL/min onto the trapping column for 6 minutes. Subsequently, peptides were eluted via the analytical column with a constant flow of 0.3 µL/min with increasing percentage of solvent B (0.1% formic acid in acetonitrile). The outlet of the analytical column was coupled directly to a QExactive plus (Thermo) mass spectrometer using the Nanospray Flex™ ion source in positive ion mode. The peptides were introduced into the QExactive plus via a Pico-Tip Emitter 360 µm OD x 20 µm ID; 10 µm tip (New Objective) and an applied spray voltage of 2.3 kV. The capillary temperature was set at 320°C. Full mass scan was acquired with mass range 375-1200 m/z in profile mode with resolution of 70000. The filling time was set at maximum of 10 ms with a limitation of 3x106 ions. Data dependent acquisition (DDA) was performed with the resolution of the Orbitrap set to 35000, with a fill time of 120 ms and a limitation of 2x105 ions. A normalized collision energy of 32 was applied. Dynamic exclusion time of 30 s was used. The peptide match algorithm was set to 'preferred' and charge exclusion 'unassigned', charge states 1, 5 - 8 were excluded. MS data was acquired in profile mode.

#### Mass Spectrometry Data Analysis

IsobarQuant[14] and Mascot (v2.2.07) were used to process the acquired data, which was searched against a Mus musculus GRCm38 pep database containing common contaminants and reversed sequences. The following modifications were included into the search parameters: Carbamidomethyl (C) and TMT10 (K) (fixed modification), Acetyl (Protein N-term), Oxidation (M) and TMT10 (N-term) (variable modifications). For the full scan (MS1) a mass error tolerance of 10 ppm and for MS/MS (MS2) spectra of 0.02 Da was set. Further parameters were set: Trypsin as protease with an allowance of maximum two missed cleavages: a minimum peptide length of seven amino acids; at least two unique peptides were required for a protein identification. The false discovery rate on peptide and protein level was set to 0.01. The raw output files of IsobarQuant (protein.txt – files) were processed using the R programming language (ISBN 3-900051-07-0). Only proteins that were quantified with at least two unique peptides were considered for the analysis. Raw TMT intensities (signal_sum columns) were first cleaned for batch effects using limma[15] and further normalized using vsn (variance stabilization normalization)[16]. Proteins were tested for differential expression using the limma package. The replicate information was added as a factor in the design matrix given as an argument to the 'lmFit' function of limma. A protein was annotated as a hit with a false discovery rate (fdr) smaller 5 % and a fold-change of at least 50 % and as a candidate with a fdr below 25 % and a fold-change of at least 50 %.

#### scRNA-seq Analysis

Processed gene count data from previously published cardiac single cell RNA-seq data[17] was downloaded from the GEO database (IDs: GSM4376680 – GSM4376710). Gene raw counts were post processed to correct gene names as well as duplicate gene entries. Seurat v4.1 was employed for single cell data processing[18] following the integration approach[19]. Subsequently, data was converted for usage with the *shinycell* web application as described by Ouyang et al[20]. All processed data is available online at: https://shiny.jakobilab.org/Hofmann_et_al_2022/.

#### Statistical Analysis

Statistical analysis was performed using GraphPad Prism 7.0 (Graphpad Software Inc; www.graphpad.com) or R (R Foundation; https://www.r-project.org). Data values are mean ± standard error of the mean (SEM). For statistical analysis one-way ANOVA with Turkey post-hoc analysis was used. When only two conditions where compared, unpaired two tailed t-test was used. $p < 0.05$ was defined as significant difference. Details of the statistical analyses of the sequencing data can be found in the respective method section. Biological replicate numbers for each figure can be found in the accompanying figure legend.

#### Supplemental Tables and Data

**Online Data I:**

Proteome Sham vs 2d IR surgery male mice

**Online Data II:**

Ribo seq Sham vs 2d IR surgery male mice DEG analysis CPM 10 cutoff

**Online Data III:**

Ribo seq Sham vs 2d IR surgery male mice DEG analysis no CPM cutoff

**Online Data IV:**

RNA seq Sham vs 2d IR surgery male mice DEG analysis CPM 10 cutoff

**Online Data V:**

RNA seq Sham vs 2d IR surgery male mice DEG analysis no CPM cutoff

#### Online Figure Legends

**Online Figure 1. eIF4E$^{S209}$ phosphorylation is controlled by ERK1/2 and not by p38 in cardiomyocytes at baseline and during ischemia or reperfusion.**

**A**, Immunoblots and quantifications of ERK1/2$^{T202/Y204}$ and HSP27$^{S82}$ phosphorylation (downstream target of p38) in response to U0126 treatment of NRCMs, n = 2. **B**, Immunoblots and quantifications of ERK1/2$^{T202/Y204}$ and HSP27$^{S82}$ phosphorylation (downstream target of p38) in response to SB202190 treatment of NRCMs, n = 2. **C**, Immunoblots and quantifications of eIF4E$^{S209}$ phosphorylation in response to U0126 or SB202190 treatment of NRCMs at baseline, after simulated ischemia (1 hour) or ischemia (3 hours) followed by reperfusion (1 hour). Ctr (control), Isch (ischemia), Rep (reperfusion), n = 2. * indicates $p<0.05$ from control. For statistical analysis one-way ANOVA with Turkey post-hoc analysis was used for **A** to **C**. $p < 0.05$ was defined as significant difference. Error bars show standard error of the mean (SEM).

**Online Figure 2. 4EGI-1 effectively suppresses eIF4F complex formation and protein synthesis in cardiomyocytes without causing cell death.**

**A** and **C**, Quantification of NRCM apoptosis by FACS analysis 24 hours after treatment with increasing doses of 4EGI-1 (**A**), n = 2. Representative FACS plots of each condition are shown below (**C**). **B**, Quantification of puromycin incorporation 6 hours after treatment with increasing doses of 4EGI-1 in NRCMs. **D**, Representative immunoblot and quantification of eIF4F complex assembly in NRCMs by immunoprecipitation of mRNA cap-binding proteins via m7GTP-coupled agarose beads at increasing timepoints after 4EGI-1 (100µM) treatment, n = 3-4. Ctr (control), * indicates $p<0.05$ from 0µM 4EGI-1 or control. For statistical analysis one-way ANOVA with Turkey post-hoc analysis was used for **A**, **B** and **D**. $p < 0.05$ was defined as significant difference. Error bars show standard error of the mean (SEM).

**Online Figure 3. Inhibition of protein synthesis by chronic rapamycin treatment after reperfusion.**

**A**, Representative puromycin immunoblot and quantification of NRCMs 24 hours and 48 hours after rapamycin (100nM) treatment after reperfusion, n = 3. The 6-hour quantification was derived from Figure 3D. Rep (reperfusion), Veh (vehicle), * indicates $p<0.05$ from time-matched vehicle. For statistical analysis one-way ANOVA with Turkey post-hoc analysis was used. $p < 0.05$ was defined as significant difference. Error bars show standard error of the mean (SEM).

**Online Figure 4. eIF4E$^{S209}$ phosphorylation does not regulate overall translation rates and does not control ischemia/reperfusion injury in vivo.**

**A**, Representative immunoblot of eIF4E after **WT-eIF4E or eIF4E$^{S209}$ phospho-dead** expression in HEK293-T cells. **B**, Quantification of puromycin incorporation in HEK293-T cells 48 hours after transfection, n = 4-6. A 6-hour treatment of 100nM Torin1 was used as a positive control. **C**, Immunoblot and quantification of eIF4E$^{S209}$ phosphorylation in mouse left ventricular lysates at increasing timepoints after i.p. injection of 10 mg/kg eFT508. **D**, Plasma Troponin T levels in male mice treated with vehicle or eFT508 (10 mg/kg) 24 hours after I/R surgery, n = 7-9. E. Left ventricular ejection fraction of male mice 2 weeks after sham or I/R surgery operated mice treated with vehicle or 10 mg/kg eFT508, n = 5-6. Ctr (control), Rep (reperfusion), Veh (vehicle). * indicates $p<0.05$ from control. For statistical analysis one-way ANOVA with Turkey post-hoc analysis was used for **A**, **B**, **C** and **E**. An unpaired two tailed t-test was used for **D**. $p < 0.05$ was defined as significant difference. Error bars show standard error of the mean (SEM).

**Online Figure 5. Activation of the mTORC1 pathway in the myocardial border zone in response to ischemia/reperfusion.**

**A**, Representative immunoblot and quantification of S6$^{S235/236}$ phosphorylation 2 or 24 hours after I/R surgery in male mice, n = 7-8. **B**, Representative phospho-S6$^{S235/236}$ immunostaining of the infarct region and border zone 2 days after Sham or I/R surgery in mice. **C**, Immunoblot and quantification of S6$^{S235/236}$ phosphorylation 28 hours after I/R surgery in male mice treated with vehicle, 2 and 6 mg/kg rapamycin or 50 mg/kg 4EGI-1, n = 3-4. Rep (reperfusion), Veh (vehicle), Rapa (rapamycin). * indicates $p<0.05$ from sham. # indicates $p<0.05$ from 28h Rep Veh. For statistical analysis one-way ANOVA with Turkey post-hoc analysis was used for **A** and **C**. $p < 0.05$ was defined as significant difference. Error bars show standard error of the mean (SEM).

**Online Figure 6. Pharmacological mTORC1 inhibtion during reperfusion protects against I/R injury and improves long-term cardiac funtion.**

**A** and **B**, Quantification of cardiac (**A**) and hepatic (**B**) S6$^{S235/236}$ phosphorylation in male mice at different timepoints after 6 mg/kg rapamycin treatment, n = 5 (control), n = 4 (6 hours), n = 3 (24 hours) as assessed by immunoblot. **C**, S6$^{S235/236}$ phosphorylation of left ventricular lysates of vehicle or 2 and 6 mg/kg rapamycin treated mice 2 days after I/R surgery assessed by immunoblot, n = 6. **D**, Quantification of myocardial puromycin incorporation in vivo 2 days after I/R surgery in response to 2 and 6 mg/kg rapamycin treatment, n = 3-6. **E**, Representative immunostaining and quantification of TUNEL$^{+}$-cells per view by immunofluorescence in sham or I/R surgery operated mice treated with vehicle or 2 and 6 mg/kg rapamycin, n = 3-4. **F**, Plasma Troponin T levels at 24 hours after I/R surgery in female (n = 3-14) and male mice (n = 4-11). **G**, 24-hour survival of all sham or I/R surgery operated mice after vehicle or 2 and 6 mg/kg rapamycin treatment used in this study, n = 31-57. **H**, 2-week survival of sham or I/R surgery operated mice treated with vehicle or 2 and 6 mg/kg rapamycin corresponding to mice used for I to L, n = 7-24. **I**, **J**, **K** and **L**, Heart weight (HW) to body weight (BW) ratio (**I**), left ventricular ejection fraction of female (n = 3-10) and male mice (n = 4-7) (**J**), infarct size (n = 2-6, male mice) (**K**), and left ventricular expression of *Nppa*, *Nppb*, *Myh7* and *Col1a1* (n = 6 for sham, n = 10 for vehicle and n = 8 for rapamycin, female mice) (**L**) 2 weeks after sham or I/R surgery operated mice treated with vehicle or 2 and 6 mg/kg rapamycin. Rep (reperfusion), Veh (vehicle), Rapa (rapamycin). * indicates p<0.05 from control or sham. # indicates p<0.05 from 2wk Rep Veh. For statistical analysis one-way ANOVA with Turkey post-hoc analysis was used for **A**, **B**, **E**, **F**, **I**, **J**, **K** and **L**. An unpaired two tailed t-test was used for **C**. A survival curve analysis was performed with GraphPad Prism 7.0 for **H**. p < 0.05 was defined as significant difference. Error bars show standard error of the mean (SEM).

**Online Figure 7. Quality control of RNA-Seq and Ribo-Seq data of sham or I/R surgery operated mice.**

**A** and **B**, Dot plot displaying the average fraction of raw sequence reads derived from rRNA, mtRNA, tRNA, snRNA and other ncRNA for RNA-Seq (**A**) and Ribo-Seq (**B**) data of left ventricular lysates derived from sham or I/R surgery operated mice. Only the 'cleaned' reads are used for subsequent data analysis. **C**, Ribo-seq read counts of all libraries showing periodic (usable), non-periodic, multi-mapped and non-aligning reads, when mapped to the mouse transcriptome. **D**, Beeswarm plot visualizing the sequenced ribosome footprint lengths across all samples. **E**, Bar plot showing the percentage of reads mapping to the coding sequence (CDS) and 5′ and 3′ untranslated regions (UTR) of annotated protein-coding genes. Each line represents a separate sample. **F**, Bar plot summarizing the ribosome protected footprint periodicity for all samples as the percentage of footprints that match the three reading frames of the annotated coding sequence genome wide. **G**, Graphical representation of the periodic profile Bayesian model selection showing results for different read lengths and p-site offsets for one typical library. **H**, Periodic footprint lengths and P-offset for all used libraries. **I** and **J**, Histograms showing the expression level of genes as measured by RNA-seq (**I**) and Ribo-Seq (**J**). Expression levels of all genes across all samples are included. Genes that met our expression cutoff of 10 counts per million (CPM) are colored red. **K** and **L**, Principal component analysis of RNA-seq (**K**) and Ribo-seq (**L**) libraries after sham or I/R surgery. Rep (reperfusion). Detailed information on statistical analysis can be found in the supplementary methods.

**Online Figure 8. Blood cell count of mice after I/R surgery and treatment with rapamycin or 4EGI-1.**

**A** to **E**, Blood cell count of red blood cells (**A**), leucocytes (**B**), lymphocytes (**C**), granulocytes (**D**) and monocytes (**E**) 24 hours after I/R surgery in male mice in treated with vehicle, 2 and 6 mg/kg rapamycin or 50 mg/kg 4EGI-1, n = 3-13. Rep (reperfusion), Veh (vehicle), Rapa (rapamycin). * indicates $p<0.05$ from pre-surgery. For statistical analysis one-way ANOVA with Turkey post-hoc analysis was used for **A** to **E**. $p < 0.05$ was defined as significant difference. Error bars show standard error of the mean (SEM).

**Online Figure 9. Bone marrow proliferative cells of mice after I/R surgery and treatment with rapamycin or 4EGI-1.**

**A** and **B.** Flow-cytometry based enumeration of bone marrow c-Kit$^+$/Sca1$^+$ cells (**A**) and bone marrow c-Kit$^+$/Sca1$^+$/Ki67$^+$ cells (**B**) 2 days after sham or I/R surgery in animals treated with vehicle, 2 and 6 mg/kg rapamycin, or 50 mg/kg 4EGI-1, n = 2-6. Rep (reperfusion), Veh (vehicle). For statistical analysis one-way ANOVA with Turkey post-hoc analysis was used. $p < 0.05$ was defined as significant difference. Error bars show standard error of the mean (SEM).

**Online Figure 10. Clustering of cardiac cells based on gene expression from sham or I/R surgery operated mice.**

**A**, Uniform manifold approximation and projection (UMAP) clustering of cardiac cells from sham

(1d and 14d) and I/R operated mice (1d, 3d, 14d). **B**, Number of cells per cluster originating from sham or I/R surgery operated mice. **C** to **F**, Violin plots showing the mRNA expression of marker genes of main cardiac cell types to identify cardiomyocytes (**C**), fibroblasts (**D**), endothelial cells (**E**) and immune cells (**F**). Data are shown as normalized transcript counts on a color-coded linear scale. Online Figures 8A to 8F were generated from publicly available mouse single-cell transcriptomics data published by Molenaar et al[17].

#### Online Figure 1

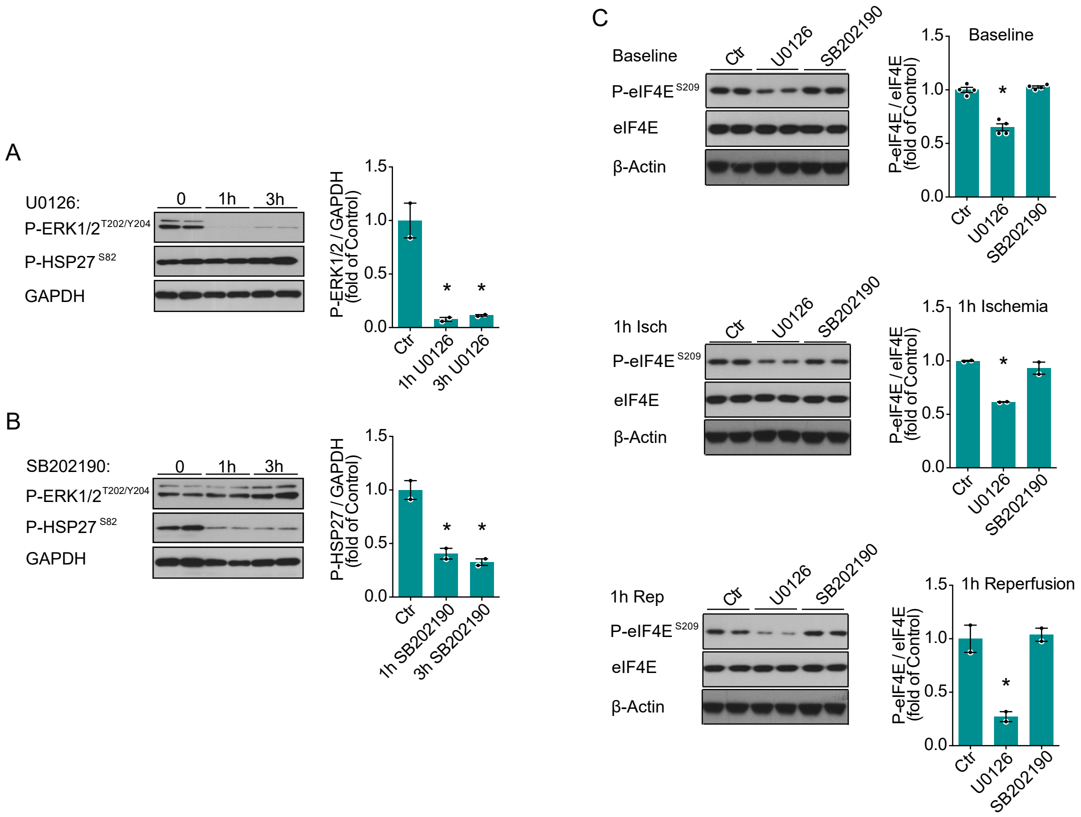

#### Online Figure 2

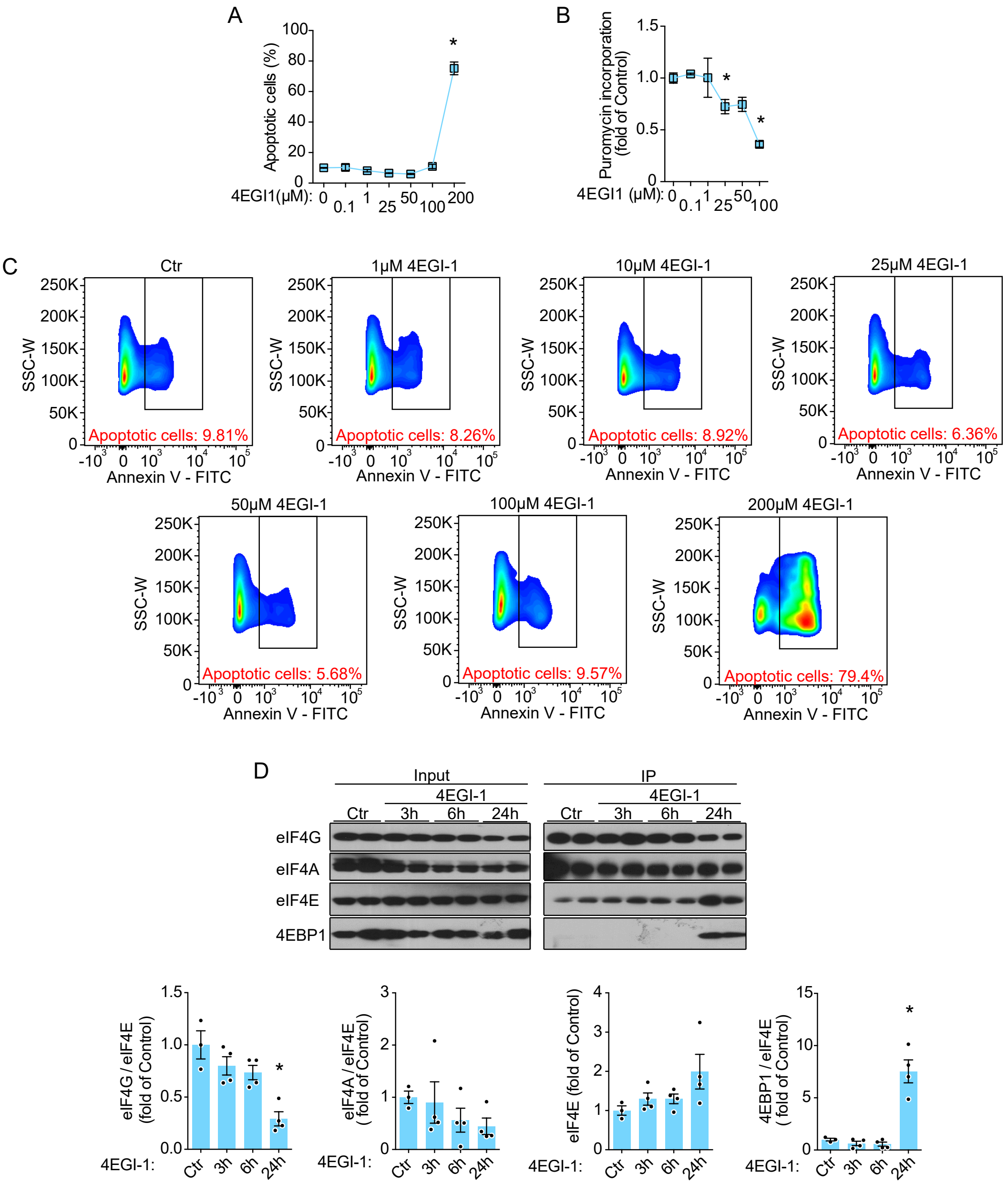

#### Online Figure 3

A

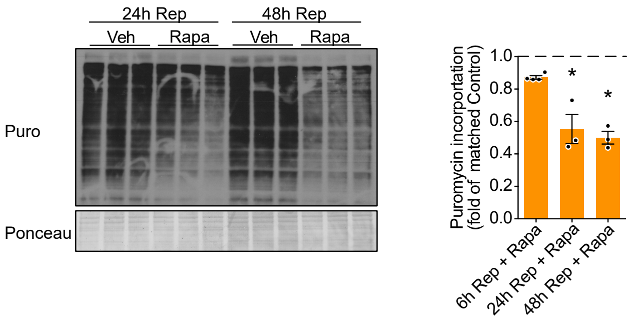

#### Online Figure 4

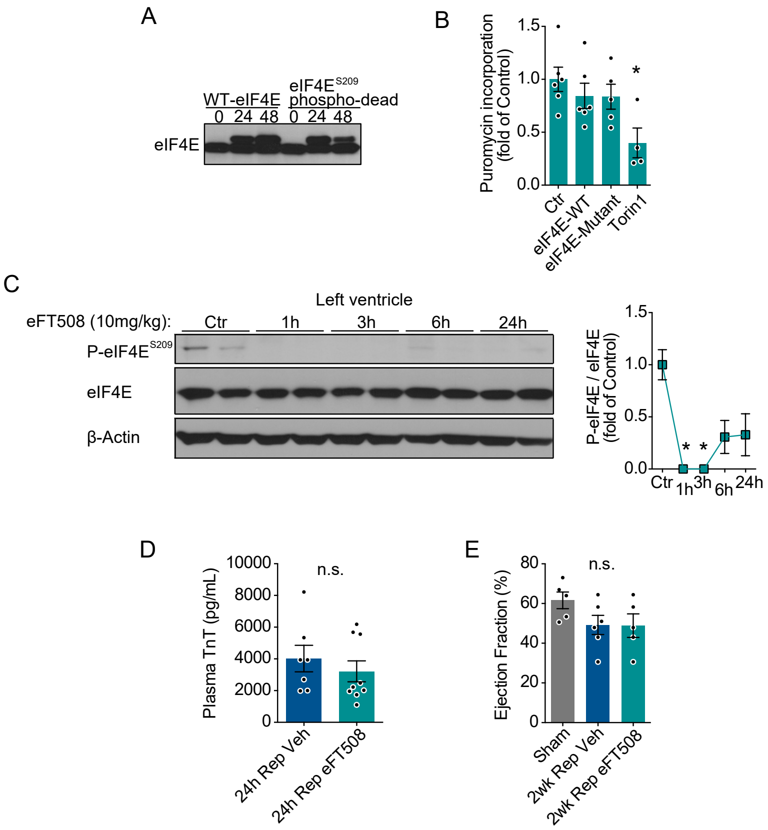

#### Online Figure 5

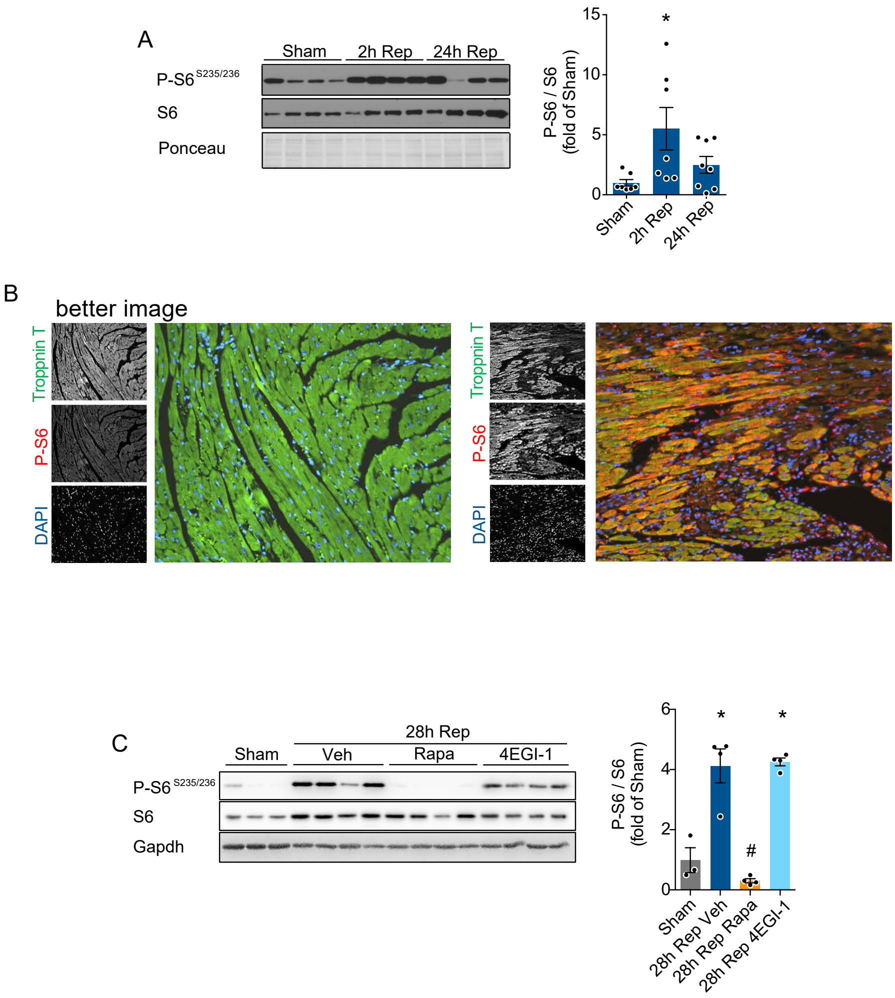

#### Online Figure 6

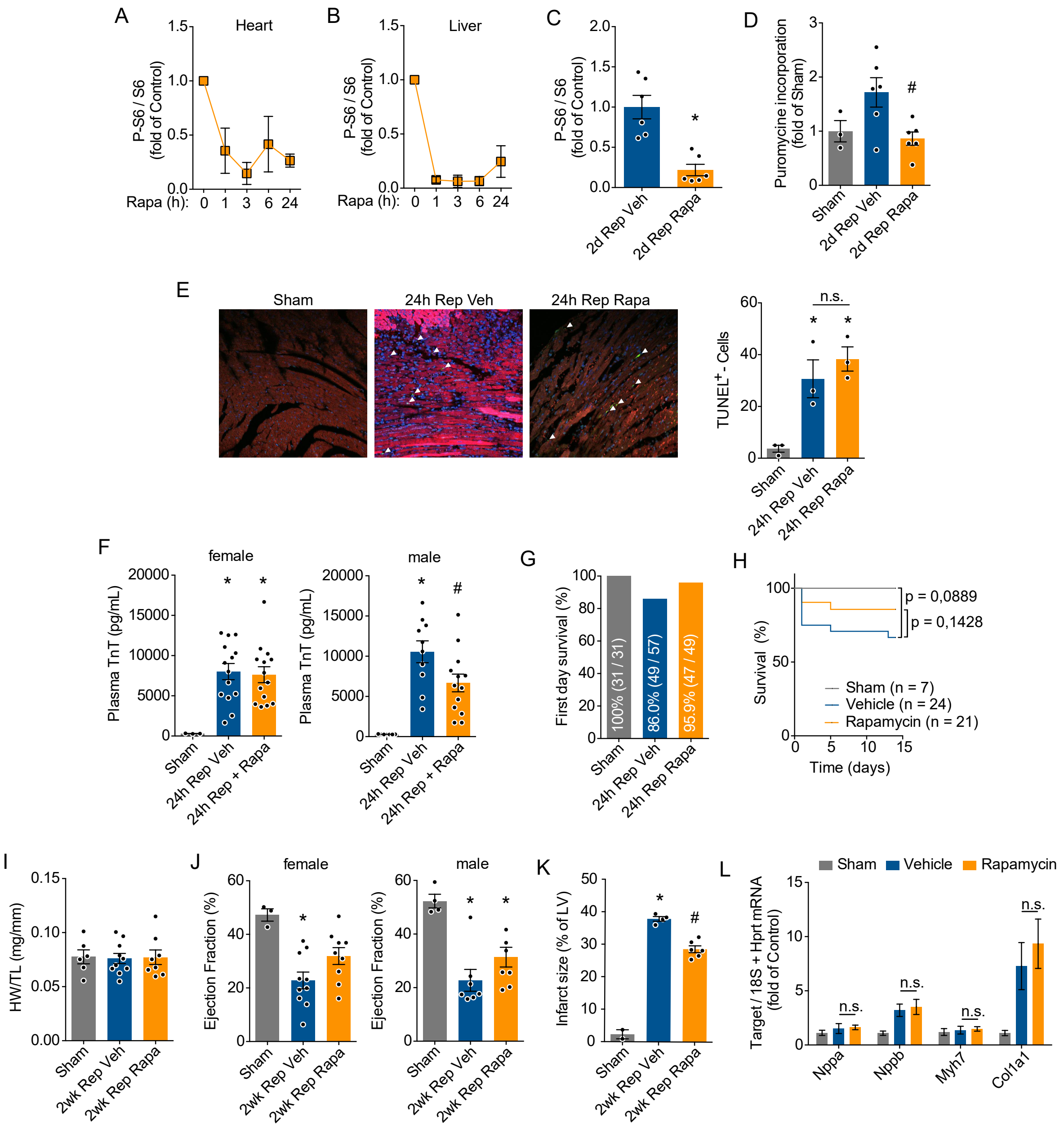

#### Online Figure 7

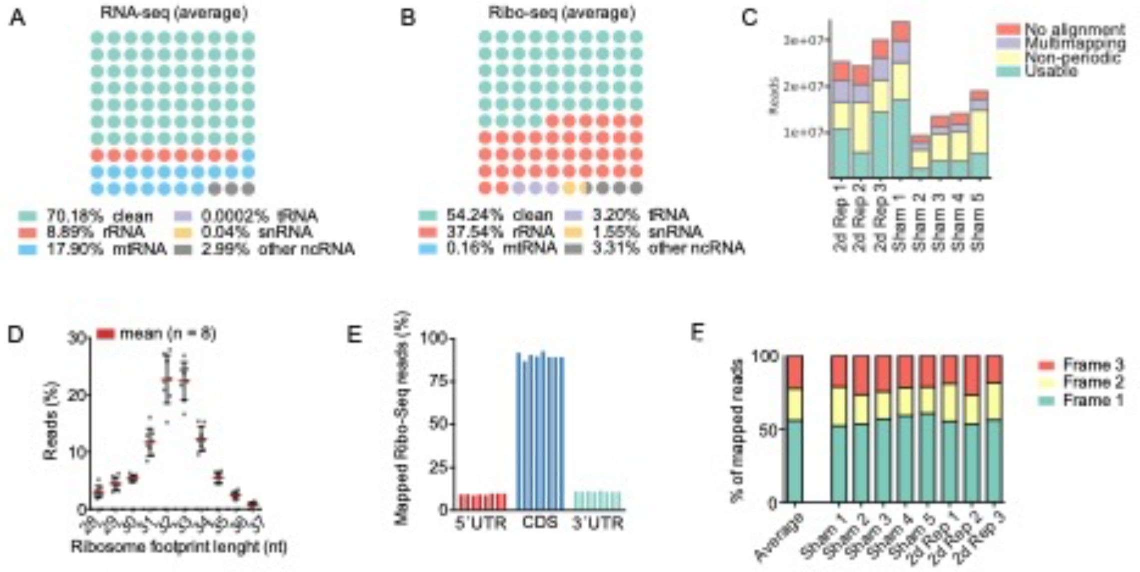


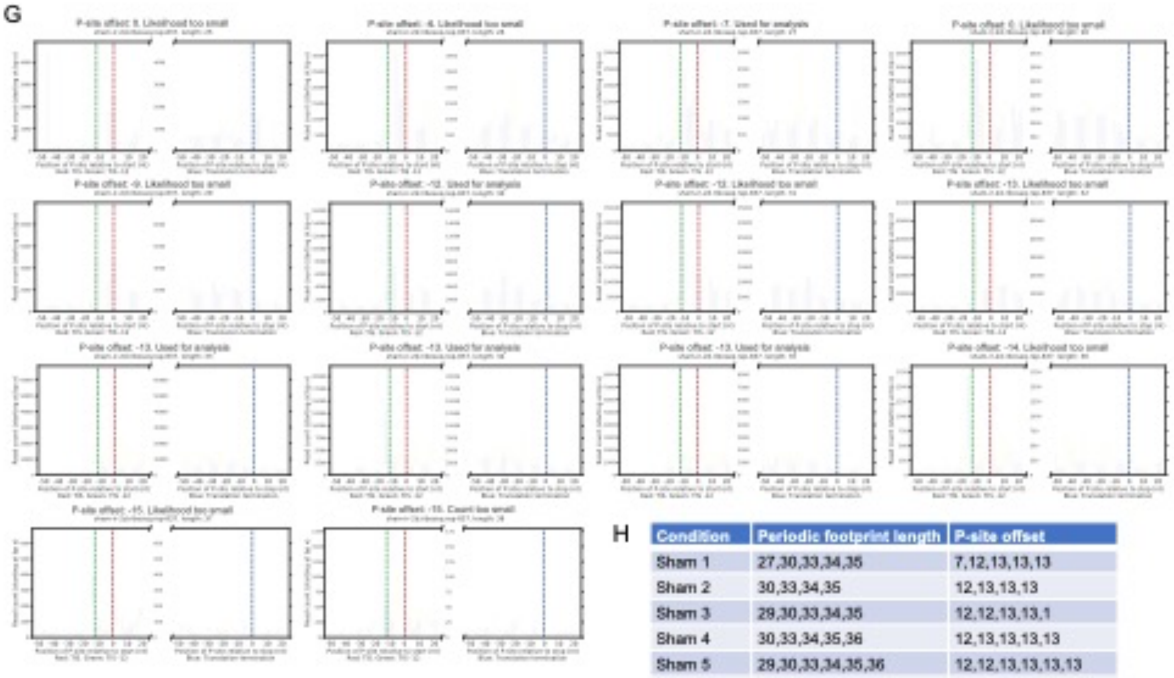


H

| Condition | Periodic footprint length | P-site offset |
|---|---|---|
| Sham 1 | 27,30,33,34,35 | 7,12,13,13,13 |
| Sham 2 | 30,33,34,35 | 12,13,13,13 |
| Sham 3 | 29,30,33,34,35 | 12,12,13,13,1 |
| Sham 4 | 30,33,34,35,36 | 12,13,13,13,13 |
| Sham 5 | 29,30,33,34,35,36 | 12,12,13,13,13,13 |
| Rep 2d 1 | 30,32,33,34,35 | 12,13,13,13,13 |
| Rep 2d 2 | 30,33,34,35 | 12,13,13,13 |
| Rep 2d 3 | 30,32,33,34,35 | 12,13,13,13,13 |

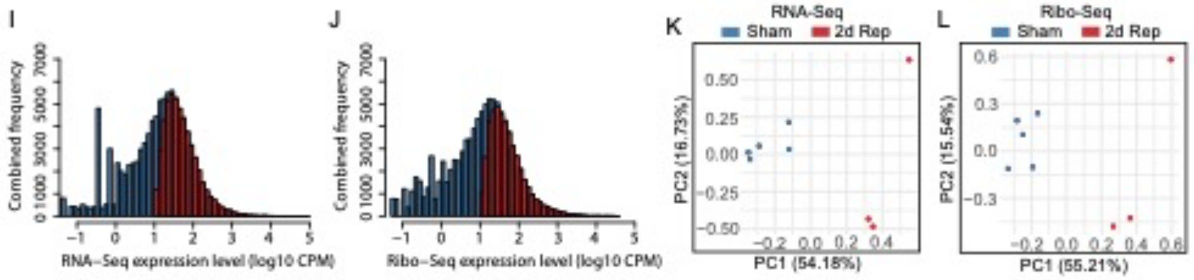

#### Online Figure 8

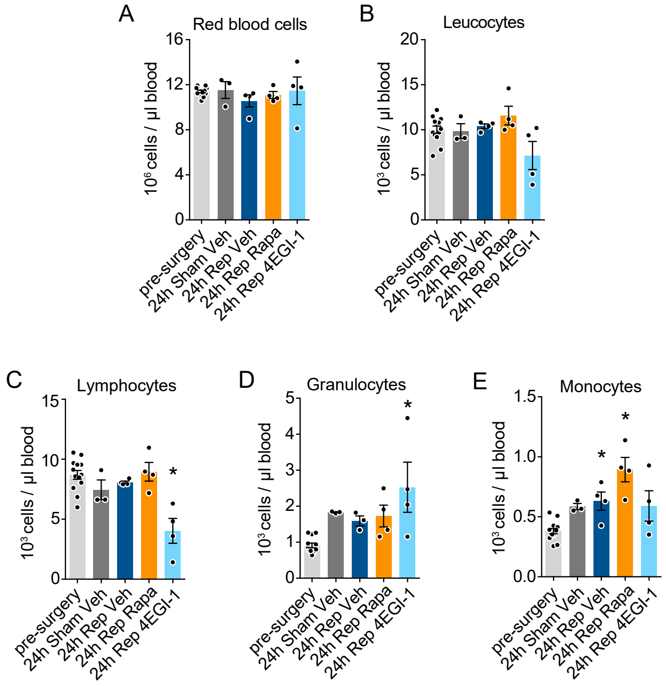

#### Online Figure 9

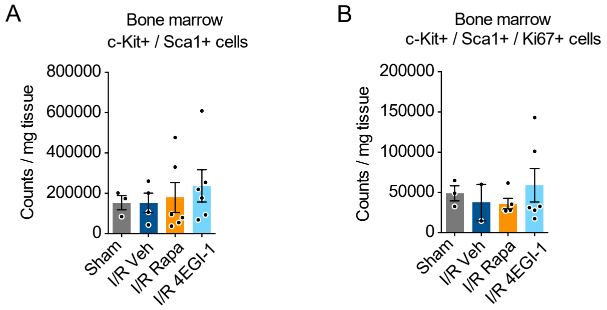

### Online Figure 10

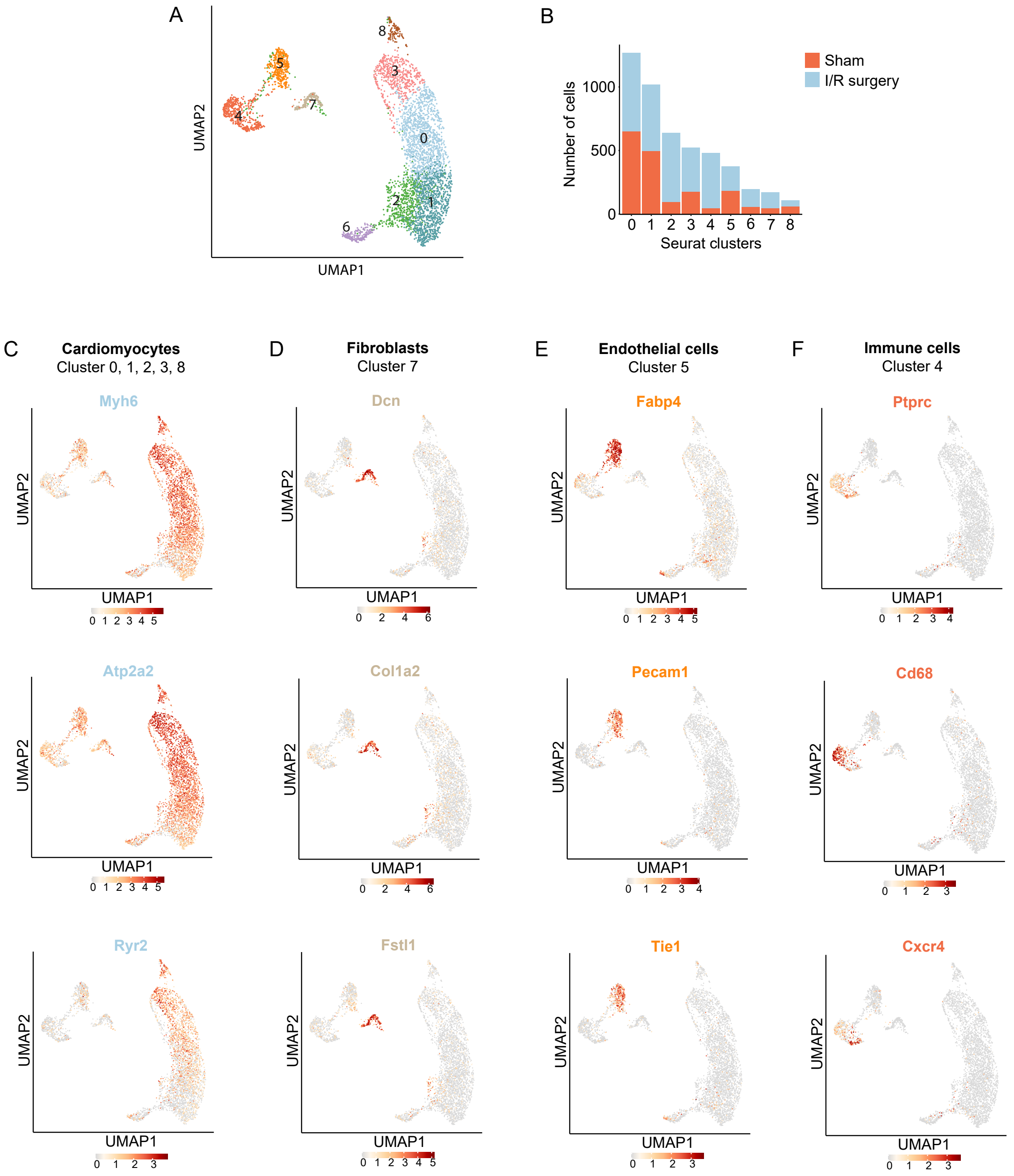
